# Cell state plasticity emerging from co-regulated, competitive, and configurable interactions within the AP-1 network

**DOI:** 10.1101/2025.11.06.686987

**Authors:** Yonatan N. Degefu, Magda Bujnowska, Douglas G. Baumann, Mohammad Fallahi-Sichani

**Affiliations:** Department of Biomedical Engineering, University of Virginia, Charlottesville, VA 22908, USA; Department of Biochemistry and Molecular Genetics, University of Virginia, Charlottesville, VA 22908, USA; Medical Scientist Training Program, University of Virginia, Charlottesville, VA 22908, USA; UVA Comprehensive Cancer Center, University of Virginia, Charlottesville, VA 22908, USA

## Abstract

AP-1 transcription factors have been implicated in cellular plasticity, differentiation-state heterogeneity, and phenotype switching in response to cancer therapies. Although AP-1 states, defined by combinatorial expression of AP-1 proteins, are heterogeneous within cell populations, only a subset of possible states is observed. How these states are constrained, why their distributions vary across cell populations, and what drives their phenotypically consequential transitions remain unclear. We develop a mechanistic ODE model of the AP-1 network, capturing dimerization-dependent, co-regulated, and competitive interactions. Calibrated to single-cell protein measurements across diverse melanoma populations and combined with statistical learning, the model reveals network features explaining population-specific AP-1 state distributions. These features correlate with MAPK signaling across tumor lines and individual cells. The model predicts and experiments validate adaptive AP-1 reconfiguration following MAPK inhibition, driving a dedifferentiated, therapy-resistant state that is attenuated through model-guided perturbations. These findings establish AP-1 as a configurable network and provide a quantitative framework for modulating AP-1 driven cell-state plasticity.

## Introduction

Cell state plasticity enables clonal tumor cells to occupy distinct transcriptional and phenotypic states and to switch between them, with major consequences for metastatic invasion, immune evasion, and resistance to therapy^1^. In melanoma, for example, tumors display transcriptional heterogeneity along a differentiation trajectory ranging from melanocytic (differentiated) to neural crest-like and mesenchymal-like (undifferentiated) states that may coexist within the same lesion or patient-derived clonal tumor line at different frequencies^2–5^; additional phenotypes, including stress-like states, have also been characterized^6^. Importantly, these states differ in their propensity to promote primary tumor growth, metastatic dissemination, and resistance to MAP kinase (MAPK)-targeted therapies^7–13^. However, despite detailed descriptions of these phenotypes, we lack mechanistic insight into what determines their presence, frequency, and co-existence, and how they vary from tumor to tumor.

AP-1 transcription factors, regulated by MAPK signaling, have been repeatedly implicated in tumor heterogeneity, cellular plasticity, and therapy resistance^14–26^. While most previous studies often treated AP-1 either as a single collective group in bioinformatic analyses or examined individual AP-1 proteins in isolation, our recent work identified the family of AP-1 proteins as a coordinated network that orchestrates melanoma cell state heterogeneity and plasticity^26^. Specifically, we showed that the relative expression of AP-1 transcription factors defines an “AP-1 state” that predicts differentiation heterogeneity across genetically diverse melanoma populations at single-cell resolution^26^. Perturbing this balance shifted the composition of cell states, implicating AP-1 as a driver rather than a mere correlate of phenotypic heterogeneity. However, the mechanisms that generate AP-1 state heterogeneity across genetically diverse tumor lines and within clonal populations, and how MAPK signaling influences these mechanisms, remain unclear.

AP-1 transcription factors function as dimeric complexes formed by bZIP-domain JUN and FOS family members that bind related DNA motifs but exhibit distinct DNA affinities and transcriptional activities depending on dimer composition^27–30^. AP-1 subunits compete for limited binding partners to form homo- and heterodimers and these dimers can auto- and cross-regulate AP-1 gene expression^31–34^. Furthermore, ERK, JNK, and p38 MAP kinases regulate basal AP-1 transcription, protein stability, and activity via transcriptional inputs and post-translational phosphorylation^35–37^. This layered architecture, including competition, feedback, and signal-mediated regulation, can potentially create a large combinatorial space from which discrete AP-1 expression states could emerge. However, we lack a mechanistic framework that integrates these regulatory principles to explain: (i) why some clonal melanoma populations are constrained to a limited number of AP-1 states, whereas others span broader AP-1 expression patterns associated with differentiation state heterogeneity; (ii) which regulatory configurations account for the distinct AP-1 compositions observed across genetically diverse populations or within heterogeneous clonal populations; (iii) what dynamic features of the AP-1 network enable AP-1 state plasticity; and (iv) how MAPK signaling shapes these configurations and transitions, before and after pathway inhibition. Addressing these gaps will reveal how AP-1 network dynamics drive non-genetic heterogeneity and plasticity in melanoma and other cellular contexts.

In this paper, we couple highly multiplexed single-cell measurements with mechanistic, single-cell ordinary differential equation (ODE) modeling and statistical learning to systematically interrogate the AP-1 network in 19 genetically distinct melanoma cell populations, representing diverse patterns of differentiation state heterogeneity. Our single-cell analysis identifies six recurrent AP-1 expression states that are non-randomly distributed within and across tumor cell populations and closely track melanoma differentiation states. We develop a mechanistic model that captures dimerization-controlled, competitive, auto- and cross-regulatory AP-1 interactions. Following calibration to single-cell data, the model explains inter- and intra-clonal AP-1 heterogeneity and links AP-1 regulatory parameters to differential MAPK signaling activities across melanoma cell populations. Through iterative experimental and model-based analyses, we reveal an adaptive reconfiguration of the AP-1 network following MAPK pathway inhibition, including JUND loss, that drives melanoma cells from a melanocytic (MITF^High^/ SOX10^High^) state to an undifferentiated (MITF^Low^/ SOX10^Low^) phenotype, a transition commonly linked to adaptive resistance to MAPK inhibitors in melanoma. This shift is mediated by FRA1 downregulation and cJUN upregulation, promoting a FRA2^High^ state that is observed in higher frequency in intrinsically undifferentiated melanoma tumors. As predicted by the model, the shift could be blocked by cJUN depletion or FRA1 overexpression. Together, these results and our integrative approach reveal network-level principles governing AP-1 driven plasticity in melanoma cells. They also provide a general framework to dissect and ultimately manipulate AP-1 network heterogeneity for therapeutic benefit.

## Results

### Melanoma cells express six recurrent AP-1 states linked to differentiation states across genetically diverse populations

We had previously used highly multiplexed immunofluorescence imaging, single-cell analysis, and machine learning to demonstrate that heterogeneity in melanoma differentiation states is associated with distinct patterns of AP-1 protein expression, including cFOS, cJUN, FRA1, FRA2, and JUND^26^. Multiplexed single-cell analysis revealed substantial variation in the expression of these AP-1 proteins both across and within melanoma cell populations derived from 19 genetically distinct cell lines, spanning a wide range of differentiation states defined by key melanocyte lineage markers SOX10 and MITF (Figures 1A, 1B, S1A). Here, we performed further analysis to show that individual cells exhibit diverse AP-1 states (defined by the combinatorial expression of the five AP-1 proteins) with variable frequencies across all cell lines. Of the 2^5^ = 32 theoretically possible states, 24 were observed experimentally but most of these states appeared at very low frequencies or were detected in only a few cell lines (Figure S1B). To focus on robust patterns, we identified six recurrent AP-1 states, defined as those present at >10% frequency (averaged across two biological replicates) in at least 3 of 19 cell lines, with minimal sensitivity to the thresholds used to classify cells as high or low AP-1 expressers (Figure S1C). We then quantified the frequency of these six states in each of the 19 cell lines (Figure 1C). While some clonal lines, such as COLO858, LOXIMVI, WM902B, and WM115, were enriched predominantly for two of the recurrent AP-1 states, others displayed a broader spectrum of AP-1 expression patterns. For instance, A375, MMACSF, and A101D exhibited high heterogeneity, with cells distributed across five or six distinct AP-1 states (Figure 1C).

**Figure 1.**
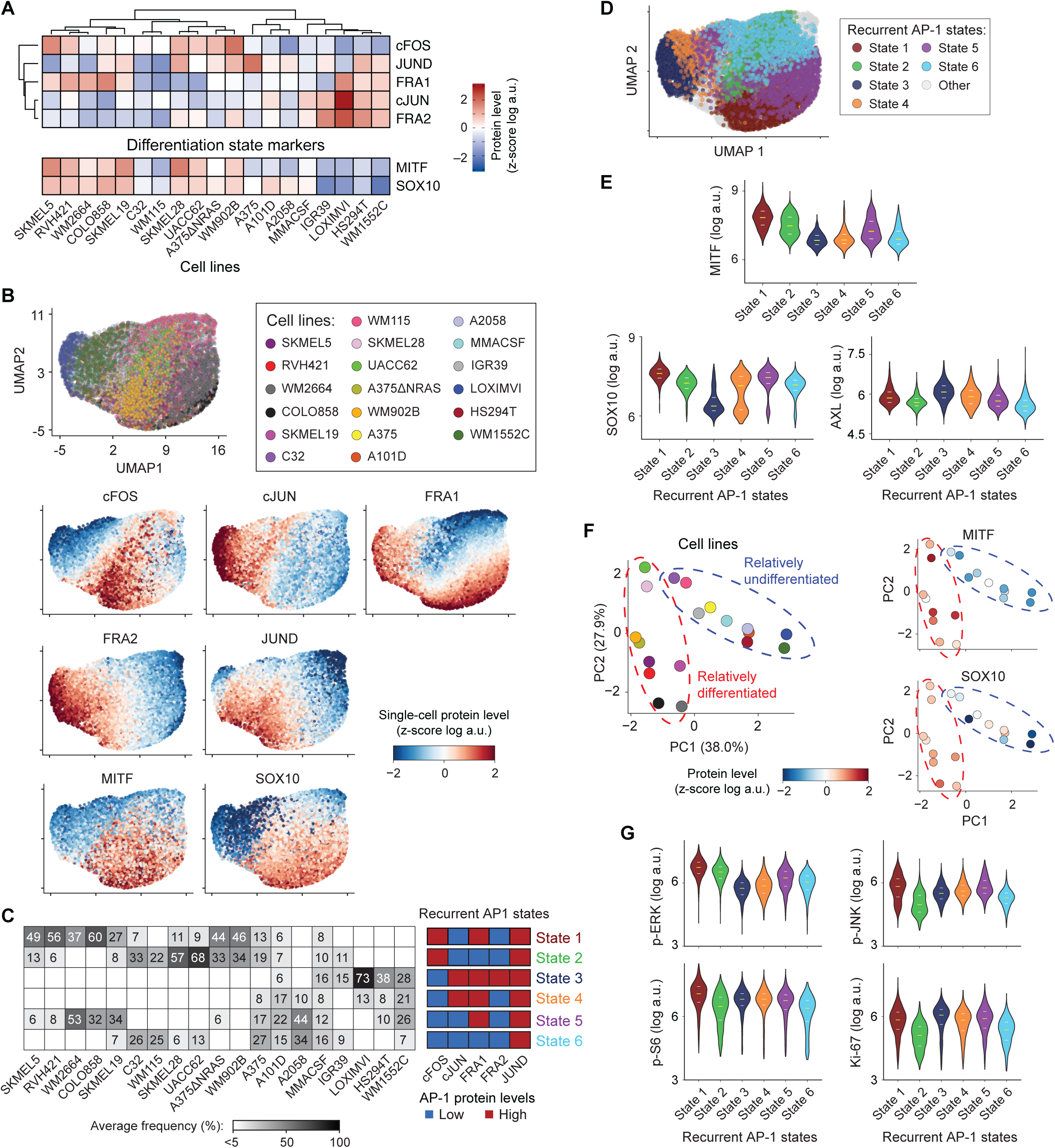
Melanoma cells express six recurrent AP-1 states linked to differentiation states across genetically diverse populations. **(A)** Population-averaged measurements of five AP-1 proteins (cFOS, FRA1, FRA2, cJUN, and JUND) and two differentiation markers (MITF and SOX10) measured by 4i across 19 BRAF-mutant melanoma cell lines. For each condition, data represent log-transformed mean values from two biological replicates, followed by z-scoring across cell lines. Data are organized by hierarchical clustering of AP-1 protein expression using Pearson correlation distance and the average linkage algorithm for computing inter-cluster distances. **(B)** UMAP visualization of melanoma cell populations based on multiplexed single-cell measurements of five AP-1 proteins (cFOS, FRA1, FRA2, cJUN, and JUND). A total of 500 randomly sampled cells per line are included in the analysis. Cells are colored either by their cell line identity (top) or by single-cell expression levels of each of the five AP-1 proteins or differentiation markers MITF and SOX10 (bottom). **(C)** Frequency of cells in each of the six recurrent AP-1 states (defined by the combinatorial expression of five AP-1 proteins) across 19 melanoma cell lines. Data are averaged across two biological replicates. **(D)** UMAP visualization of melanoma cells as in (B) colored by AP-1 state. **(E)** Single-cell distributions of differentiation markers MITF, SOX10 and AXL protein levels among the six recurrent AP-1 states combined across all cell lines. Violin plots show median and interquartile ranges. **(F)** Principal component analysis (PCA) of melanoma cell lines based on their single-cell frequencies in the six recurrent AP-1 states shown in (C). Cell lines are colored either by identity (left) or by population-averaged expression of differentiation markers (MITF and SOX10) (right). **(G)** Single-cell distributions of p-ERK^T202/Y204^, p-JNK^T183/Y185^, p-S6^S235/S236^, and Ki-67 protein levels among the six recurrent AP-1 states combined across all cell lines. Violin plots show median and interquartile ranges.

Single-cell analysis illustrated how the six recurrent AP-1 states were distributed across the diverse panel of cell lines (Figure 1D). It also revealed how these states differed in the abundance of AP-1 proteins and melanoma differentiation markers, measured in the same cells by iterative indirect immunofluorescence imaging (4i) (Figures S1D, 1D-1E). For example, AP-1 state 1 (cFOS^High^/ cJUN^Low^/ FRA1^High^/ FRA2^Low^/ JUND^High^) was associated with the highest levels of both MITF and SOX10, transcription factors co-expressed in melanocytic (differentiated) cells, whereas AP-1 state 3 (cFOS^Low^/ cJUN^High^/ FRA1^High^/ FRA2^High^/ JUND^High^) showed the lowest levels of these markers and instead expressed the highest levels of AXL, consistent with an undifferentiated phenotype (Figure 1E). Principal component analysis (PCA) further grouped cell lines by the diversity and frequency of the six recurrent AP-1 states, revealing an unsupervised correlation with their population-averaged differentiation states (Figure 1F). Cell lines enriched for differentiated (MITF^High^/ SOX10^High^) cells, such as COLO858 and SKMEL5, were separated from predominantly undifferentiated (MITF^Low^/ SOX10^Low^) lines such as LOXIMVI, IGR39 and HS294T (Figure 1F), and this separation corresponded with differences in cFOS, FRA2, and cJUN levels (Figure S1E).

In addition to their correlation with melanoma differentiation state markers, the recurrent AP-1 states differed in the phosphorylation of MAP kinases (p-ERK^T202/Y204^ and p-JNK^T183/Y185^), phosphorylation of ribosomal protein S6 (p-S6^S235/S236^; an integrator of RAS/RAF/MAPK signaling), and the expression of the proliferation marker Ki-67 (Figure 1G). For example, AP-1 states 1 and 2, both distinguished by high cFOS expression, displayed highest levels of p-ERK; and AP-1 states 2 and 6, which share low levels of cJUN, FRA1, and FRA2, exhibited the lowest levels of p-JNK, p-S6, and Ki-67 (Figure 1G).

Together, these results confirm our previous findings that AP-1 expression is closely associated with melanoma differentiation state heterogeneity at a single-cell level. Furthermore, they reveal that a subset of six distinct and recurrent AP-1 expression states, each associated with differences in differentiation markers and MAPK pathway activities, is distributed in non-random patterns across genetically diverse melanoma cell populations. However, what drives the heterogeneity of AP-1 states across and within clonal cell lines, and what might govern the co-existence of specific states in some clonal populations but not in others, remains unclear. Importantly, the apparent cell-to-cell differences captured in static snapshots of AP-1 states in a clonal population may reflect dynamic transitions between these states. To explore this possibility and uncover the underlying determinants of AP-1 state heterogeneity, we used computational modeling to examine the dynamic features of AP-1 network interactions that could give rise to the observed heterogeneity across cell populations.

### Computational modeling shows dimerization-controlled competitive regulation drives single-cell AP-1 heterogeneity

To build a mechanistic computational model of AP-1 network dynamics in melanoma cells, we focused on key network features that are likely to underlie the emergence of heterogeneous AP-1 expression states. A key feature of AP-1 is the ability of its subunits to form a range of homo- and heterodimers via conserved bZIP domains. Different dimer combinations exhibit distinct DNA-binding affinities and transcriptional activities^27–30^. Because individual JUN and FOS family proteins can participate in multiple dimeric configurations, the relative abundance of these components gives rise to context-dependent competition for binding partners, introducing combinatorial complexity that influences which AP-1 dimers predominate in a given cellular state. In addition, AP-1 proteins differ in stability and can regulate their own and each other’s expression in a dimer-specific manner^38–40^. These auto- and cross-regulatory interactions can amplify expression differences and stabilize specific AP-1 states over time^41^. We thus developed a computational model to test whether these core mechanisms can explain the observed diversity of AP-1 states and to identify key dynamic features that shape AP-1 state distributions within and across clonal melanoma cell populations. To this end, we assembled a minimal set of rules based on literature evidence describing interactions among five key AP-1 proteins: cFOS (encoded by the *FOS* gene), FRA1 (encoded by *FOSL1*), FRA2 (encoded by *FOSL2*), cJUN (encoded by *JUN*), and JUND (encoded by *JUND*)

The model represents the dynamics of AP-1 protein dimerization, production, and degradation in individual cells using ordinary differential equations (ODEs) (Figure 2A; See Methods for additional details). AP-1 dimerization is governed by protein-specific association (*k_on_*) and dissociation (*k_off_*) rate constants. JUN subfamily proteins (cJUN, JUND) can form homodimers, while FOS family members (cFOS, FRA1, FRA2) require a JUN partner to dimerize. Each AP-1 protein (X) is produced at a basal rate (α*_X_*) representing its constitutive expression. For proteins subject to auto- or cross-regulation, an additional dimer-induced production term (β*_X_by_YZ_*) captured their regulated expression, modeled as a Hill function of the concentration of the dimer (YZ) that induces their transcription. These include cJUN expression induced by cJUN-cJUN and cJUN-cFOS dimers, FRA1 expression induced by cJUN-FRA1 and JUND-cFOS, and FRA2 expression induced by cJUN-FRA2^30,40,42–46^. Four dimers not involved in AP-1 auto- or cross-regulation included JUND-JUND, JUND-cJUN, JUND-FRA1, JUND-FRA2^30,40,47,48^. Each AP-1 protein degrades at a rate (γ) that depends on its identity and dimerization state. We used the ODE model to track the concentrations of all AP-1 proteins (including monomers and dimers) at the single-cell level. To enable comparison with experimental data, we calculated the total concentration of each of the five AP-1 subunits (cFOS, cJUN, FRA1, FRA2, and JUND), regardless of their dimerization state.

**Figure 2.**
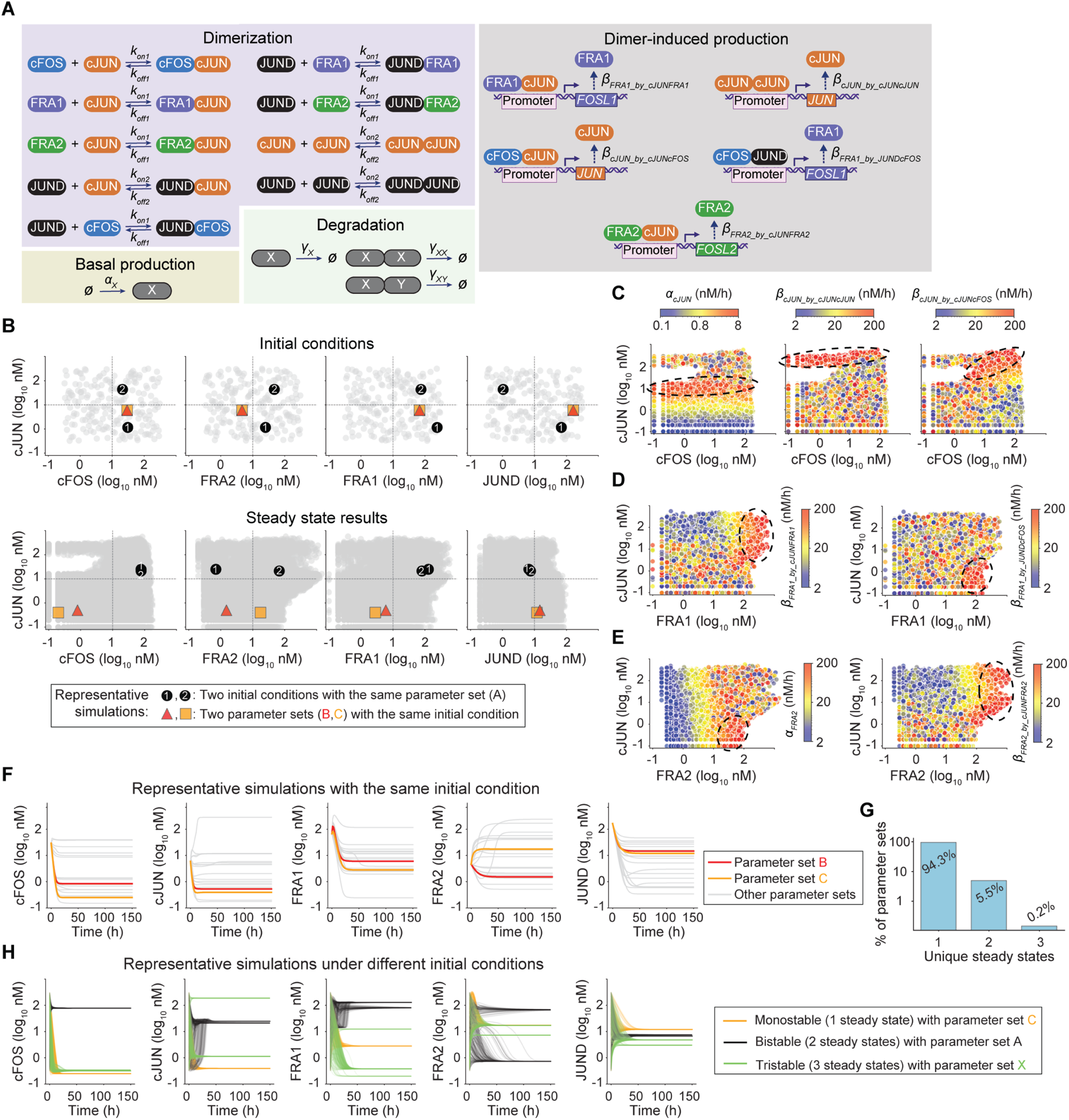
Computational modeling shows dimerization-controlled competitive regulation drives single-cell AP-1 heterogeneity. **(A)** Schematic illustration of reactions and associated rate constants governing AP-1 network dynamics, including protein dimerization, basal production, dimer-induced production, and degradation in individual cells. **(B)** Visualization of AP-1 initial conditions (top) and steady state outcomes (bottom) from 4 million dynamic simulations, combining 200 initial conditions with 20,000 parameter sets sampled by LHS. Representative simulations are highlighted: black circles 1 and 2 indicate runs with distinct initial conditions but the same parameter set A, yielding different steady states due to system bistability; the red triangle and orange square represent simulations with the same initial condition, but distinct parameter sets B and C, producing divergent steady states. **(C-E)** Visualization of AP-1 steady state outcomes as a function of individual parameters representing basal or dimer-induced production of cJUN (C), FRA1 (D), and FRA2 (E). Dashed circles highlight enriched outcomes associated with the highest values of the indicated parameters. **(F)** Time-course dynamics of representative simulations with identical initial conditions but distinct parameter sets. Simulations associated with parameter sets B (red) and C (orange) correspond to those highlighted in panel (B). **(G)** Percentage of parameter sets (out of 20,000 sampled by LHS) that converged to a single steady state, bistability, or tristability. **(H)** Time-course dynamics of representative simulations resulting in a single steady state (parameter set C, orange), two steady states (parameter set A, black), or three steady states (parameter set X, green).

To explore potential sources of cell-specific heterogeneity in AP-1 states, we first performed a global parameter scan using Latin Hypercube Sampling (LHS), an efficient approach for sampling multiple parameters simultaneously from a multidimensional distribution^49^. We varied two classes of cell-specific inputs: (i) the initial concentrations of the five AP-1 proteins and (ii) 15 kinetic parameters representing basal production, dimer-induced production, and degradation rates for each AP-1 protein. Kinetic parameters, including fixed rate constants for dimerization and dissociation, as well as estimated ranges for protein production and degradation, were informed by prior studies in related mammalian systems (Table S1). Simulations were run to steady state using a 1000-fold range of initial AP-1 concentrations, spanning from very low (0.316 nM) to very high (316 nM) levels. In total, we performed 4 million simulations by combining 200 initial conditions with 20,000 randomly sampled parameter sets.

Visualization of steady state results from all simulations revealed the theoretical limits of AP-1 expression, constrained by the structure of AP-1 network interactions across the full 15-dimensional parameter space (Figure 2B). For example, total cJUN expression varied with all three rate parameters that control its production, including its basal production (α*_cJUN_*) and induced production by cJUN-cJUN and cJUN-cFOS dimers (β*_cJUN_by_cJUNcJUN_*and β*_cJUN_by_cJUNcFOS_*, respectively) (Figures 2C, S2A-S2C). However, highest cJUN expression levels (>100 nM) were achieved when induced production via dimer-mediated feedback was active. In contrast, maximal basal production alone resulted in only moderately high cJUN levels (10-20 nM). A similar pattern was observed for FRA1, where total expression levels >100 nM were achieved under either high β*_FRA1_by_cJUNFRA1_* or high β*_FRA1_by_JUNDcFOS_* values (Figures 2D, S2D, S2E). The former corresponded to cJUN^High^/ FRA1^High^ states, while the latter was predominantly associated with cJUN^Low^/ FRA1^High^ states. Likewise, varying basal FRA2 production (α*_FRA2_*) versus dimer-induced production by cJUN-FRA2 (β*_FRA2_by_cJUNFRA2_*) enriched for distinct AP-1 states with respect to cJUN and FRA2 levels (Figures 2E, S2F and S2G). Collectively, these parameters have strong control over AP-1 dynamics, with their variation giving rise to diverse AP-1 expression states (Figure 2F).

A hallmark of nonlinear regulatory systems with feedback, competition, and cooperativity, all of which are embedded in the structure of AP-1 network, is multi-stability^50^. In such systems, differences in initial protein levels can drive cells toward alternative steady states, enabling cell state heterogeneity even under identical parameter conditions. To evaluate whether the AP-1 network supports multi-stability within physiologically plausible parameter ranges, we analyzed the 20,000 parameter sets sampled by LHS, each tested across 200 distinct initial conditions also chosen by LHS. Among all simulations, 94.3% of parameter sets resulted in convergence to a single steady state regardless of initial conditions, while 5.5% exhibited bistability and 0.2% displayed tristability (Figures 2G and 2H). These results indicate that while most parameter combinations support a unique AP-1 expression state, a subset of specific regulatory configurations allows for multiple stable outcomes, consistent with the idea that AP-1 network architecture can intrinsically drive diversification of cell states. However, not all theoretically possible configurations are observed experimentally. To identify the subset of model outcomes that could explain AP-1 state heterogeneity across melanoma cell populations, we next calibrated the parameter space to single-cell data from each of the 19 cell lines.

### Differences in calibrated AP-1 parameters explain AP-1 state diversity across genetically distinct melanoma populations

To identify regulatory configurations consistent with experimentally observed AP-1 state distributions, we calibrated the model separately for each of the 19 melanoma cell lines. To enable direct comparison with experimental data (measured by quantitative imaging in arbitrary units), we discretized total protein levels in the simulations using a threshold of 10 nM: expression levels ≥10 nM were considered high, and <10 nM were considered low. This binarization allowed us to classify each simulation into one of the six experimentally defined AP-1 expression states and systematically identify parameter sets compatible with the heterogeneity observed in each cell line.

Since each cell line exhibits a distinct subset of the six recurrent AP-1 states, we filtered simulations to retain only those parameter sets and initial conditions that reproduced one or more of the observed states as steady state outcomes, either through retention (cells remaining in a given state) or transition (cells moving between observed states) (Figure 3A). For example, for COLO858 which is composed of only two states (cFOS^High^/ cJUN^Low^/ FRA1^High^/ FRA2^Low^/ JUND^High^, and cFOS^Low^/ cJUN^Low^/ FRA1^High^/ FRA2^Low^/ JUND^High^), we selected simulations in which an initial state either remained stable or transitioned to the other (Figure 3A). For cell lines composed of three states, a broader set of simulations was retained, including those that captured any of the six possible transitions among the three states, as well as stable retention of each state. Cell lines exhibiting all six recurrent AP-1 states (such as MMACSF and A101D) were compatible with the largest subset of simulation outcomes, encompassing the full spectrum of possible state transitions.

**Figure 3.**
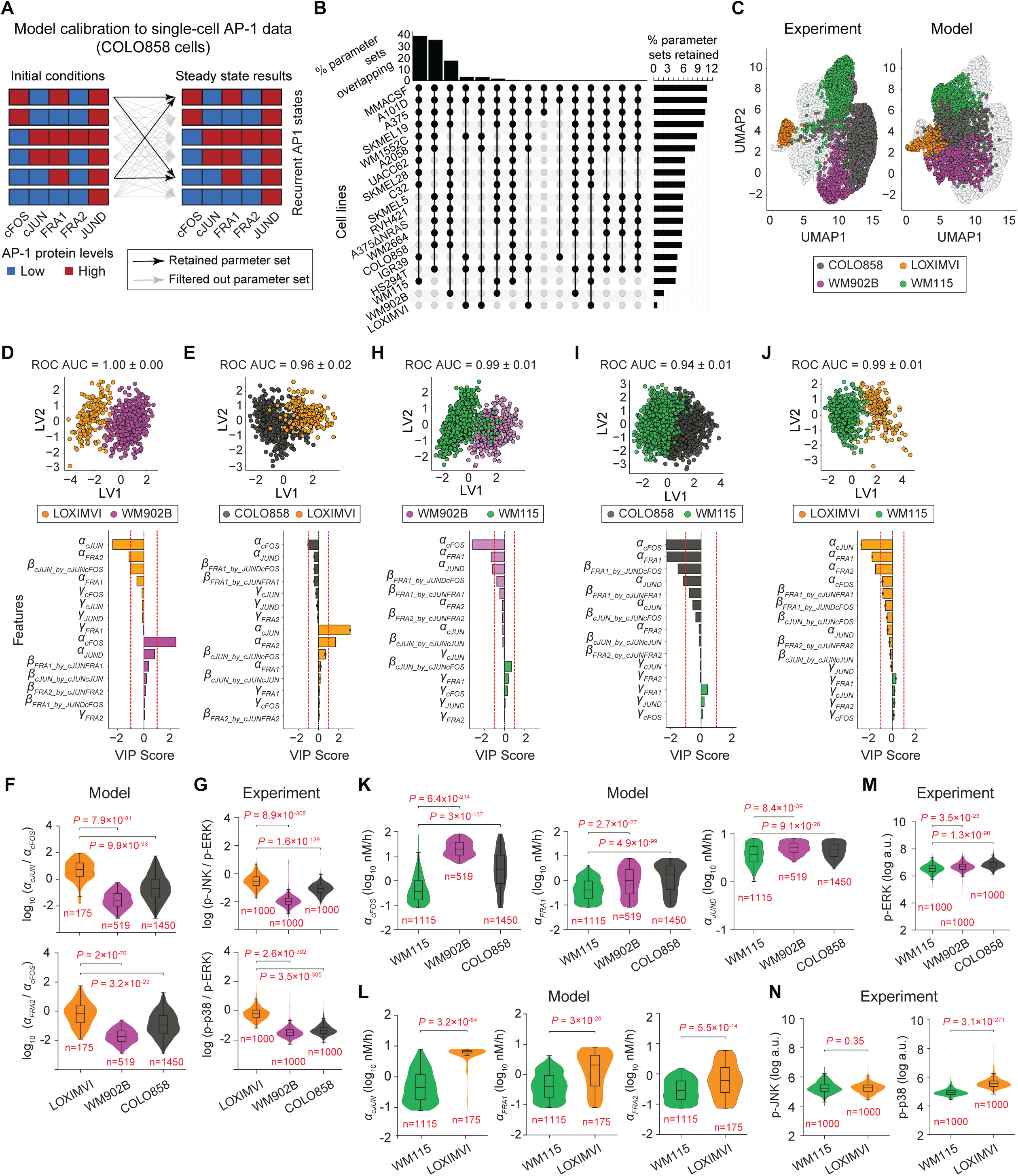
Differences in calibrated AP-1 parameters explain AP-1 state diversity across genetically distinct melanoma populations. **(A)** Schematic illustration of ODE model calibration to single-cell multiplexed AP-1 data. For each cell line, simulations were filtered to retain only parameter sets and initial conditions that reproduced one or more experimentally observed AP-1 states as steady state outcomes, either through retention (cells remaining in a given state) or transition (cells switching between observed states). Arrows indicate all theoretically possible transitions among the six recurrent AP-1 states; bold arrows highlight retained simulations for the example cell line COLO858. **(B)** Experimentally plausible parameter configurations for each cell line (bar plots on the right) and groups of shared regulatory configurations across cell lines (bar plots on top) obtained after model calibration to single-cell data from each line. **(C)** Visualization of z-scored steady state AP-1 concentration values from the ODE model (right) alongside z-scored single-cell experimental data (left) for four melanoma cell lines (COLO858, LOXIMVI, WM902B, and WM115) projected onto a shared UMAP embedding. **(D-E)** PLS-DA models comparing cell populations from LOXIMVI versus WM902B (D) and COLO858 versus LOXIMVI (E) using the 15 calibrated model parameters as predictive features. PLS-DA score plots (LV1 versus LV2) show individual cells colored by cell line identity (top), and signed VIP scores identify top discriminating parameters (bottom). Model performance was evaluated by five-fold stratified cross-validation, with the area under the ROC curve (AUC) reported as mean ± SD. **(F)** Model-predicted single-cell distributions of the α*_cJUN_*/α*_cFOS_* ratio (top) and α*_FRA2_*/α*_cFOS_* ratio (bottom) among three melanoma cell lines: LOXIMVI, WM902B, and COLO858. **(G)** Experimentally measured single-cell distributions of p-JNK/p-ERK (top) and p-p38/p-ERK (bottom) among the same cell lines. **(H-J)** PLS-DA models comparing cell populations from WM902B versus WM115 (H), COLO858 versus WM115 (I), and LOXIMVI versus WM115 (J). PLS-DA score plots display individual cells colored by cell-line identity (top) and signed VIP scores identify the top discriminating parameters (bottom). **(K-L)** Model-predicted single-cell distributions of α*_cFOS_*, α*_FRA1_* and α*_JUND_* among WM115, WM902B, and COLO858 (K) and of α*_cJUN_*, α*_FRA1_* and α*_FRA2_* between WM115 and LOXIMVI (L). **(M-N)** Experimentally measured single-cell distributions of p-ERK among WM115, WM902B, and COLO858 cell lines (M) and of p-JNK and p-p38 between WM115 and LOXIMVI (N). Statistical comparisons in (F), (G), and (K-N) were performed using two-sided Wilcoxon rank-sum tests. Box plots show medians and interquartile ranges.

Model calibration substantially reduced the parameter space, narrowing it down to experimentally plausible configurations for each cell line (Figure 3B; see rows). The extent of reduction varied depending on the diversity of AP-1 states observed in each line. For example, fewer than 1% of all simulations were retained for LOXIMVI, the most tightly constrained cell line, which exhibited only two AP-1 states (cFOS^Low^/ cJUN^High^/ FRA1^High^/ FRA2^Low^/ JUND^High^, and cFOS^Low^/ cJUN^High^/ FRA1^High^/ FRA2^High^/ JUND^High^). In contrast, over 10% of simulations were retained for MMACSF and A101D, two highly heterogeneous cell lines that exhibited all six recurrent AP-1 states. In addition to substantially narrowing the parameter space, the analysis revealed shared regulatory configurations across cell lines by grouping the calibrated parameter sets into 15 subgroups based on their compatibility with overlapping AP-1 state compositions (Figure 3B; see columns). Comparison of such experimentally plausible parameter configurations across cell lines provides an opportunity to uncover potential molecular mechanisms underlying differences in AP-1 state distributions across distinct clonal populations. To test this idea, we focused on relatively well-calibrated melanoma cell lines (e.g., LOXIMVI, WM902B, WM115 and COLO858) which represent distinct differentiation states and exhibit markedly different patterns of AP-1 state distribution. To determine whether the calibrated computational models for these cell lines capture differences in experimentally observed AP-1 state distribution, we combined z-scored steady state AP-1 concentration values from the ODE model with z-scored single-cell experimental data from all four cell lines and projected them onto a shared UMAP embedding. Simulated single-cell AP-1 states for each cell line clustered closely with their corresponding experimental counterparts (Figure 3C). Although some AP-1 states are shared between cell lines, contributing to partial overlap in UMAP space, each cell line is composed of a unique combination of AP-1 states that distinguishes it from the others. For example, WM902B shares one AP-1 state with COLO858 (cFOS^High^/ cJUN^Low^/ FRA1^High^/ FRA2^Low^/ JUND^High^) and another with WM115 (cFOS^High^/ cJUN^Low^/ FRA1^Low^/ FRA2^Low^/ JUND^High^), resulting in partial overlap with those cell lines. In contrast, both AP-1 states observed in LOXIMVI are unique to this cell line, consistent with its greater separation from the other three populations in UMAP space (Figure 3C).

To identify the molecular parameters that may drive these AP-1 state differences, we constructed partial least squares discriminant analysis (PLS-DA) models comparing pairs of cell lines using the underlying model parameters as predictive features. For example, a model comparing LOXIMVI (an undifferentiated cell line) and WM902B (a differentiated cell line) discriminated between the two cell populations with high accuracy (ROC AUC ≈ 1; Figures 3D and S3A). Variable importance in projection (VIP) analysis identified a higher basal production rate of cFOS (α*_cFOS_*) in WM902B and higher basal production rates of cJUN (α*_cJUN_*) and FRA2 (α*_FRA2_*) in LOXIMVI as the top contributors to their distinct AP-1 state compositions (Figure 3D; bottom). A similar PLS-DA model comparing LOXIMVI with COLO858 (another differentiated cell line) also achieved high classification accuracy (ROC AUC ≈ 0.96; Figures 3E and S3B) and identified the same three parameters (α*_cFOS_*, α*_cJUN_* and α*_FRA2_*) as key determinants of AP-1 state differences between the two lines (Figure 3E; bottom).

The basal expression of AP-1 proteins is regulated by multiple signaling pathways, including MAP kinases ERK, JNK, and p38^40^. Among these, JUND is expressed constitutively and its expression is regulated by sustained ERK signaling, especially under mitogenic stimulation^51^. In BRAF-mutant melanomas, where ERK signaling is constitutively hyperactivated, this leads to uniformly high JUND expression across cells^26^. In contrast, cFOS and cJUN are classic immediate early genes whose expression is more transient and tightly regulated primarily by mitogenic ERK and stress-responsive JNK signaling pathways, respectively^40^. FRA1 and FRA2 are also inducible by MAPK pathways, but their regulation is more delayed and context-dependent, typically requiring sustained input^52^. While both are primarily regulated by ERK, FRA2 is additionally co-regulated by p38 and JNK signaling, particularly under stress conditions^53^. Given these differences in upstream regulation, we asked whether the distinct basal production rates of cFOS, cJUN, and FRA2 predicted by the model in BRAF-mutant LOXIMVI, WM902B, and COLO858 cells might reflect differences in the relative activity of MAPK signaling pathways. To test this, we quantified ERK, JNK, and p38 activity at the single-cell level using multiplexed immunofluorescence imaging of their phosphorylated forms: p-ERK^T202/Y204^, p-JNK^T183/Y185^ and p-p38^T180/Y182^, respectively. Consistent with model predictions showing higher α*_cJUN_*/α*_cFOS_* and α*_FRA2_*/α*_cFOS_* ratios in LOXIMVI cells compared to WM902B and COLO858 (Figure 3F), we observed correspondingly elevated p-JNK/p-ERK and p-p38/p-ERK ratios in LOXIMVI relative to both WM902B and COLO858 (Figure 3G).

Similar findings emerged from PLS-DA comparisons involving WM115, another relatively undifferentiated line, when analyzed against other cell lines. The basal production rates of cFOS, FRA1, and JUND were predicted to be lower in WM115 compared to the differentiated lines WM902B and COLO858 (Figures 3H, 3I, S3C and S3D), while the dominant differences between LOXIMVI and WM115 were predicted to result from higher production rates of cJUN, FRA1, and FRA2 in LOXIMVI (Figures 3J and S3E). Consistent with these predictions (Figures 3K and 3L), we experimentally observed elevated p-ERK levels in WM902B and COLO858 relative to WM115 (Figure 3M), and elevated p-p38 levels in LOXIMVI compared to WM115 (Figure 3N). Together, these results show that the calibrated model can identify parameter configurations that are consistent with the diversity of AP-1 states observed across genetically distinct melanoma cell populations. Differences in inferred AP-1 regulatory parameters are associated with cell line-specific MAPK activity, supporting a link between differential upstream signaling and the emergence of distinct AP-1 expression states.

### Variation in AP-1 production and stability drives intra-clonal AP-1 state heterogeneity consistent with MAPK signaling differences

In addition to differences in AP-1 states observed across genetically distinct melanoma cell lines, each clonal cell line also exhibits cell-to-cell heterogeneity, with individual cells occupying at least two distinct AP-1 states. While variability among cell lines may reflect underlying genetic differences, heterogeneity within a clonal population is likely driven by non-genetic mechanisms. We therefore asked which molecular parameters might underlie this non-genetic AP-1 state heterogeneity within clonal populations. To address this, we constructed PLS-DA models to compare pairs of recurrent AP-1 states co-existing within each clonal cell line. In COLO858 cells, cells shared a cJUN^Low^/ FRA1^High^/ FRA2^Low^/ JUND^High^ profile but varied in cFOS expression (cFOS^Low^ versus cFOS^High^). WM902B cells were cFOS^High^/ cJUN^Low^/ FRA2^Low^/ JUND^High^ with a mixture of FRA1^Low^ and FRA1^High^ profiles. LOXIMVI cells were cFOS^Low^/ cJUN^High^/ FRA1^High^/ JUND^High^ but exhibited variation in FRA2 expression (FRA2^Low^ versus FRA2^High^). We thus used PLS-DA to identify model parameters distinguishing these intra-line AP-1 states: cFOS^Low^ versus cFOS^High^ in COLO858, FRA1^Low^ versus FRA1^High^ in WM902B, and FRA2^Low^ versus FRA2^High^ in LOXIMVI.

In COLO858, the PLS-DA model identified the basal production rate of cFOS (α*_cFOS_*) as the most significant feature distinguishing cFOS^Low^ and cFOS^High^ subpopulations (Figures 4A and S4A). Consistent with this prediction, cFOS^Low^ cells exhibited significantly lower levels of p-ERK than cFOS^High^ cells (Figure 4B). In WM902B cells, the model identified several parameters enriched in the FRA1^High^ subpopulation: a higher basal production rate of FRA1 (α*_FRA1_*), higher induced production of FRA1 by the JUND-cFOS dimer (β*_FRA1_by_JUNDcFOS_*), and greater molecular stability of FRA1 (i.e., lower degradation rate, γ*_FRA1_*) (Figures 4C and S4B). In line with no predicted difference in α*_cFOS_*, the two subpopulations showed no significant difference in p-ERK levels (Figure 4D; top panel). However, FRA1^High^ cells showed elevated Serine 265 phosphorylation of FRA1 (p-FRA1^S265^), known to enhance FRA1 protein stability (Figure 4D; middle panel). The FRA1^High^ subpopulation also showed higher JUND protein levels compared to FRA1^Low^ cells, consistent with the predicted role of JUND-cFOS dimer in FRA1 production (Figure 4D; bottom panel). In LOXIMVI cells, the PLS-DA model predicted the FRA2^High^ subpopulation to be associated with elevated basal production rates of FRA2 (α*_FRA2_*) and higher induced production of FRA2 by the cJUN-FRA2 dimer (β*_FRA2_by_cJUNFRA2_*) (Figures 4E and S4C). Although the FRA2^High^ and FRA2^Low^ subpopulations showed no significant differences in p-ERK or p-JNK levels (consistent with their comparable α*_cFOS_* and α*_cJUN_* values), FRA2^High^ cells exhibited significantly higher p-p38 and cJUN levels, consistent with elevated α*_FRA2_* and β*_FRA2_by_cJUNFRA2_*levels (Figure 4F).

**Figure 4.**
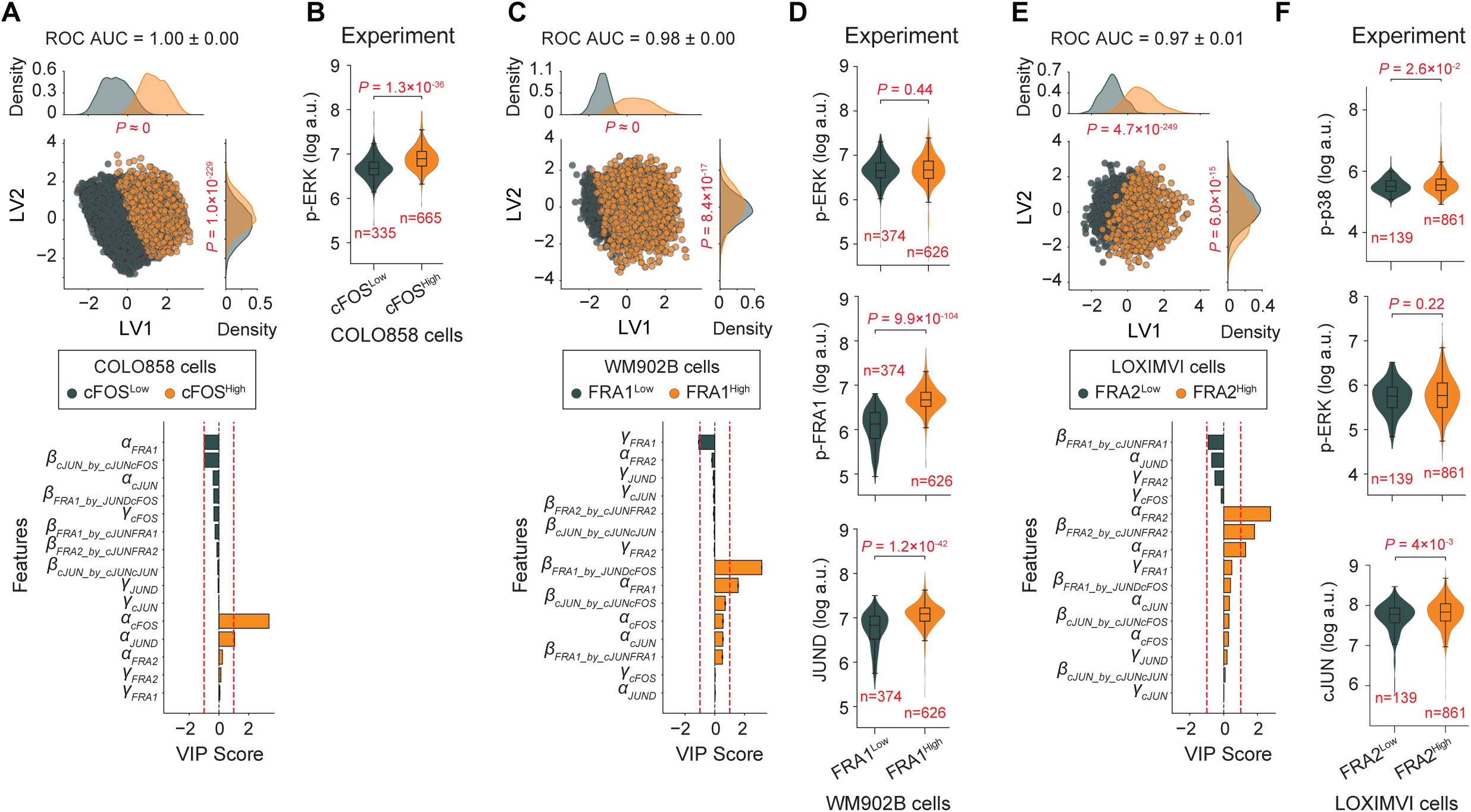
Variation in AP-1 production and stability drives intra-clonal AP-1 state heterogeneity consistent with MAPK signaling differences. **(A)** PLS-DA model comparing cFOS^High^ and cFOS^Low^ subpopulations in COLO858. PLS-DA score plot (top) shows individual cells colored by cFOS expression state, with their distributions compared across latent variables. Signed VIP scores (bottom) identify the top discriminating parameters. **(B)** Experimentally measured single-cell distribution of p-ERK between cFOS^High^ and cFOS^Low^ subpopulations in COLO858. **(C)** PLS-DA model comparing FRA1^High^ and FRA1^Low^ subpopulations in WM902B. **(D)** Experimentally measured single-cell distributions of p-ERK (top), p-FRA1 (middle), and JUND (bottom) between FRA1^High^ and FRA1^Low^ subpopulations in WM902B. **(E)** PLS-DA model comparing FRA2^High^ and FRA2^Low^ subpopulations in LOXIMVI. **(F)** Experimentally measured single-cell distributions of p-p38 (top), p-ERK (middle), and cJUN (bottom) between FRA2^High^ and FRA2^Low^ subpopulations in LOXIMVI. The performance of PLS-DA models was evaluated using five-fold stratified cross-validation, and the area under the ROC curve (AUC) is reported as mean ± SD. Statistical significance of single-cell distribution differences along each PLS-DA latent variable in (A), (C), and (E) was evaluated using Kolmogorov-Smirnov (KS) tests. Statistical comparisons in (B), (D), and (F) were performed using two-sided Wilcoxon rank-sum tests. Box plots display medians and interquartile ranges.

Together, these results highlight how variation in key regulatory parameters, including those controlling basal production, induced expression, and protein stability, can account for heterogeneity in AP-1 state composition within clonal melanoma populations, in a manner consistent with experimentally observed differences in upstream MAP kinase signaling activity.

### The balance between basal and dimer-induced FRA2 production determines FRA2 expression bistability

As noted above, most parameter combinations within physiologically plausible ranges converged to a single steady state, corresponding to a unique AP-1 expression profile. However, a subset of configurations supported bistability or even tristability, suggesting that the AP-1 network can drive diversification of cell states and enable heterogeneity within clonal populations under fixed kinetic parameter conditions. To identify parameters distinguishing monostable from bistable or tristable regimes across all cells, we constructed PLS-DA models while accounting for class imbalance via repeated random down-sampling. The models distinguished monostable regime from bistable or tristable regimes with mean ROC AUC values of 0.74 ± 0.02 and 0.92 ± 0.05, respectively (Figures 5A-5D and S5A-S5B). VIP analysis showed that both bistable and tristable outcomes were associated with substantially increased production of cJUN induced via the cJUN-cJUN dimer (β*_cJUN_by_cJUNcJUN_*) (Figures 5B and 5D).

**Figure 5.**
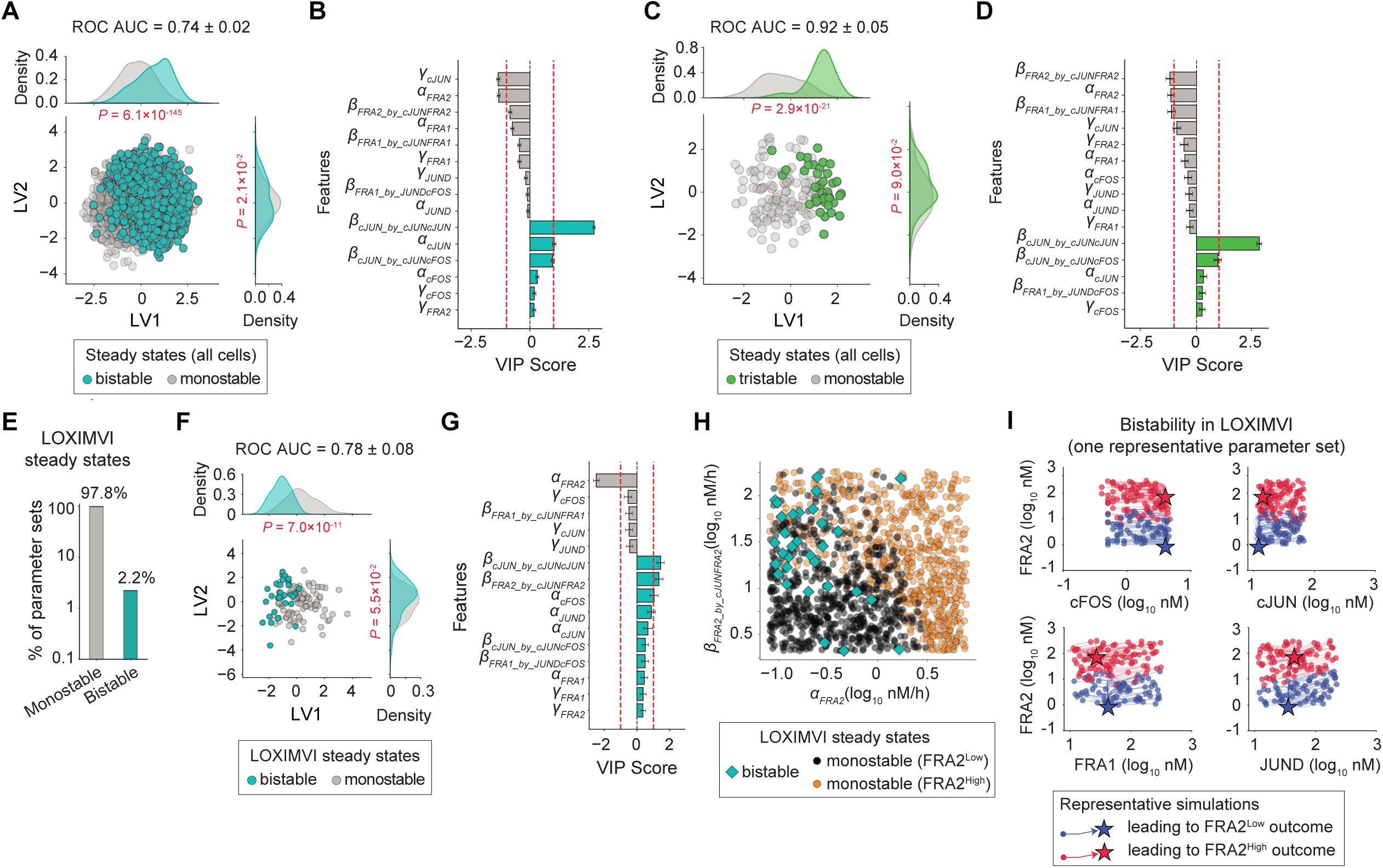
The balance between basal and dimer-induced FRA2 production determines FRA2 expression bistability. **(A-D)** PLS-DA models comparing bistable vs monostable (A,B), and tristable vs monostable (C,D) regimes across all cells. PLS-DA score plots (A,C) highlight the respective stability regimes with distributions compared across latent variables. Model performance was evaluated using five-fold stratified cross-validation, and the area under the ROC curve (AUC) is reported as mean ± SD. Statistical significance of distribution differences along each latent variable was assessed using the Kolmogorov-Smirnov (KS) test. Signed VIP scores (B,D) identify top discriminating parameters for the respective stability regime comparisons. **(E)** Percentage of parameter sets converging to a single steady state or exhibiting bistable FRA2^High^ and FRA2^Low^ states in LOXIMVI. **(F)** PLS-DA model comparing bistable and monostable regimes in LOXIMVI. The PLS-DA score plot shows individual simulations colored according to whether they represent monostable or bistable steady states, with their distributions compared across latent variables. Model performance was evaluated in a similar way as described for (A) and (C). **(G)** Signed VIP scores identify the top discriminating parameters between monostable and bistable steady states in LOXIMVI. **(H)** Distribution of bistable versus monostable (FRA2^High^ and FRA2^Low^) steady states as a function of basal FRA2 production rate (α*_FRA2_*) and cJUN-FRA2 dimer-induced FRA2 production (β*_FRA2_by_cJUNFRA2_*). **(I)** Representative simulations illustrating bistable FRA2^High^ and FRA2^Low^ states in LOXIMVI.

Among cell lines exhibiting only two of the recurrent AP-1 states, LOXIMVI was unique in that both states could arise from bistability (Figure 5E). These bistable solutions gave rise to both FRA2^High^ and FRA2^Low^ states under identical parameter settings. To identify parameters distinguishing monostable from bistable regimes in LOXIMVI, we constructed a PLS-DA model that classified these regimes with a mean ROC AUC = 0.78 ± 0.08 (Figures 5F and S5C). VIP analysis showed that monostable outcomes in LOXIMVI were associated with higher basal production of FRA2 (α*_FRA2_*), whereas bistable outcomes were characterized by both elevated β*_cJUN_by_cJUNcJUN_*(consistent with the global analysis above) as well as increased dimer-induced production of FRA2 via the cJUN-FRA2 complex (β*_FRA2_by_cJUNFRA2_*) (Figures 5G and S5D). These results suggest that the relative balance between basal and dimer-induced FRA2 production may play a role in determining FRA2 expression states in LOXIMVI. Specifically, low levels of both basal and induced production favored a monostable FRA2^Low^ state (Figure 5H; gray circles), whereas high basal production favored a monostable FRA2^High^ state regardless of induced production rates (Figure 5H; orange circles). In contrast, high dimer-induced FRA2 production combined with low basal production increased the likelihood of bistability, allowing co-existence of both FRA2^Low^ and FRA2^High^ states (Figure 5H; cyan diamonds). In this bistable regime, cells with identical rate parameters can stably occupy either state, but sufficiently large perturbations in AP-1 protein levels, without changes in underlying parameters, can trigger transitions between states (Figure 5I). Although this mechanism remains to be experimentally validated, it offers a hypothesis for how nonlinear feedback and dynamic regulation within the AP-1 network may contribute to non-genetic heterogeneity in melanoma cell populations.

### Loss of JUND following ERK inhibition induces a FRA2^High^ state and promotes transition to an undifferentiated phenotype via decreased FRA1 and increased cJUN expression

The FRA2^High^ state, which co-occurs with elevated cJUN expression, is strongly associated with the undifferentiated (SOX10^Low^/ MITF^Low^) phenotype, as demonstrated in our prior analyses of human melanoma cell lines and tumors^26^. The undifferentiated phenotype is frequently linked to increased invasiveness and intrinsic resistance to MAPK pathway inhibitors^54^. Consistent with our multiplexed protein-level analysis by 4i, the analysis of single-cell RNA-seq data from a melanoma patient-derived xenograft (PDX) model^11^ also showed a negative correlation between FOSL2 transcript abundance and the melanocytic gene signature defined by Tsoi et al.^2^ (Figure S6A), as well as positive correlations between FOSL2 and JUN transcript levels and the undifferentiated gene signature, including markers such as NGFR and AXL (Figures S6B-S6D). Although single-cell RNA-seq is limited in its ability to resolve multiplexed AP-1 network states due to transcript sparsity, this univariate analysis provides orthogonal support for the association between elevated FOSL2 and JUN levels and the undifferentiated phenotype in melanoma cells.

In the panel of melanoma cell lines profiled by 4i-based protein analysis, the FRA2^High^/ cJUN^High^ state was predominant in intrinsically undifferentiated lines such as HS294T, LOXIMVI, and IGR39, comprising approximately ∼50-90% of the cell populations (Figure S6E). Following 24 hours of ERK pathway inhibition (via treatment with the RAF inhibitor vemurafenib or in combination with the MEK inhibitor trametinib), these cell lines remained predominantly in the FRA2^High^/ cJUN^High^ state (Figure S6E). Under the same conditions, several FRA2^Low^ cell lines (including COLO858, RVH421, WM2664, and A375) upregulated FRA2 together with cJUN, resulting in a 4-60 fold expansion of the FRA2^High^/ cJUN^High^ subpopulation, from ∼0.2-3% to ∼7-26% (Figure 6A). In contrast, another subset of relatively FRA2^Low^ cell lines (including SKMEL5 and SKMEL28) failed to induce FRA2 and instead exhibited up to a 10-fold reduction in the FRA2^High^/ cJUN^High^ fraction, decreasing from ∼4-9% to ∼0.3-0.5% (Figure 6B). These treatment-induced changes in FRA2 and cJUN levels was accompanied by corresponding shifts in differentiation state, as indicated by decreased and increased MITF expression within 72 hours, respectively (Figure 6C).

**Figure 6.**
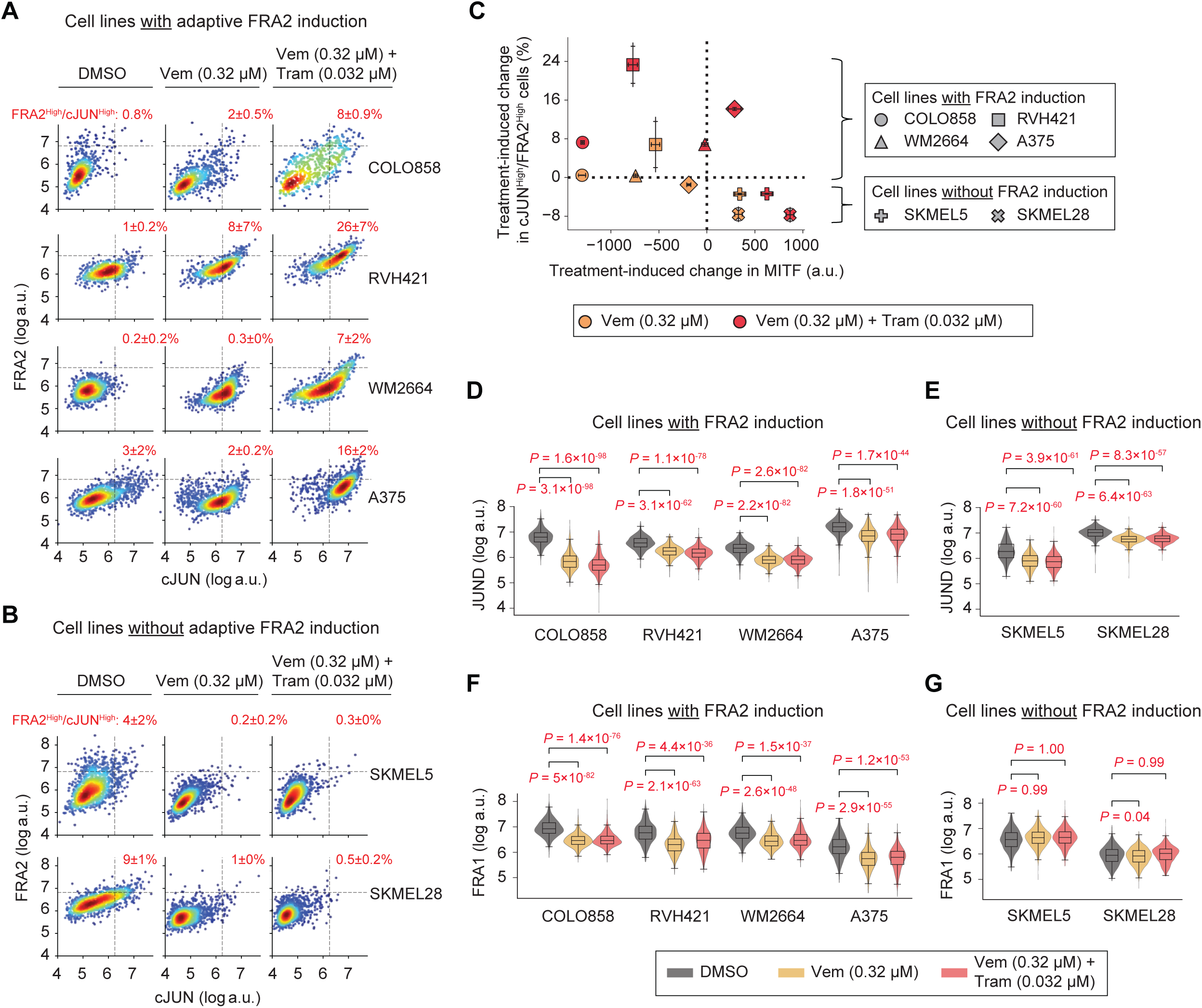
MAPK inhibitor-induced FRA2 upregulation is accompanied by cJUN upregulation, JUND downregulation, and FRA1 downregulation. **(A-B)** Single-cell covariance analysis of FRA2 and cJUN protein levels in melanoma cells following 24 h treatment with Vemurafenib (0.32 μM), the combination of Vemurafenib (0.32 μM) and Trametinib (0.032 μM), or vehicle control (DMSO), measured by 4i. Data are shown for cJUN^Low^/ FRA2^Low^ cell lines that exhibit adaptive treatment-induced upregulation of cJUN and FRA2 (A) and cJUN^Low^/ FRA2^Low^ cell lines that fail to induce either factor following MAPK inhibitor treatment (B). The percentage of cJUN^High^/ FRA2^High^ cells (averaged across two replicates ± SD) is shown in the top-right quadrant of each plot. **(C)** Population-level analysis of treatment-induced changes in the percentage of cJUN^High^/ FRA2^High^ cells (mean ± SD across two replicates) at 24 hours, and their association with changes in MITF levels at 72 hours, for the two subgroups of cell lines with and without FRA2 induction, as described in (A) and (B), respectively. **(D-G)** Single-cell analysis of JUND (D,E) and FRA1 protein levels (F,G) in melanoma cells following 24 h treatment with Vemurafenib (0.32 μM), the combination of Vemurafenib (0.32 μM) and Trametinib (0.032 μM), or DMSO, measured by 4i. Data are shown for cell lines with adaptive FRA2 induction (D and F) and FRA2^Low^ lines that do not induce FRA2 expression upon MAPK inhibitor treatment (E and G). Statistical comparisons were performed using one-sided Wilcoxon signed-rank tests. Box plots display medians and interquartile ranges.

These observations indicate that ERK pathway inhibition can induce an adaptive transition toward a FRA2^High^/ cJUN^High^ state associated with dedifferentiation in some, but not all FRA2^Low^ melanoma populations. Exploring AP-1 dynamics underlying this adaptive response, we found that JUND levels declined following ERK pathway inhibition in all of the FRA2^Low^ cell lines that induced cJUN and FRA2 (Figure 6D), but not in intrinsically cJUN^High^/ FRA2^High^ cell lines (HS294T, LOXIMVI, IGR39) (Figure S6F). However, JUND downregulation was also observed in FRA2^Low^ cell lines that failed to induce cJUN or FRA2 expression (Figure 6E). These data suggest that while JUND loss may be associated with adaptive FRA2 induction following ERK pathway inhibition, it is not predictive on its own. We thus explored whether additional factors might distinguish FRA2-inducing versus non-inducing cell lines. Notably, both FRA2-inducing cell lines and intrinsically FRA2^High^ cell lines exhibited a reduction in FRA1 expression following ERK inhibition, while non-inducing FRA2^Low^ lines did not (Figures 6F, 6G, S6G). Therefore, what distinguished FRA2-inducing cell lines from both intrinsically FRA2^High^ and non-inducing FRA2^Low^ lines was the coordinated downregulation of both JUND and FRA1 and upregulation of cJUN following ERK pathway inhibition. We thus hypothesized that the transition from a FRA2^Low^ to FRA2^High^ state, and the associated shift toward an undifferentiated phenotype, may be initiated by JUND loss but requires additional AP-1 network remodeling, including reduced FRA1 and increased cJUN, to enable this transition.

To test this hypothesis, we depleted JUND expression in COLO858 cells using both shRNA-mediated knockdown (JUND^KD^) and CRISPR/Cas9-mediated knockout (JUND^KO^). In both cases, a subpopulation of FRA2^High^ cells emerged, accompanied by downregulation of FRA1 and upregulation of cJUN, with no significant change in cFOS expression (Figure 7A). We then assessed whether this AP-1 network reconfiguration was sufficient to induce a shift in differentiation state by co-staining SOX10 and MITF. Single-cell analysis revealed a significant increase in the fraction of SOX10^Low^ /MITF^Low^ cells under both JUND^KD^ and JUND^KO^ conditions compared to controls (Figure 7B), consistent with a shift toward an undifferentiated phenotype.

**Figure 7.**
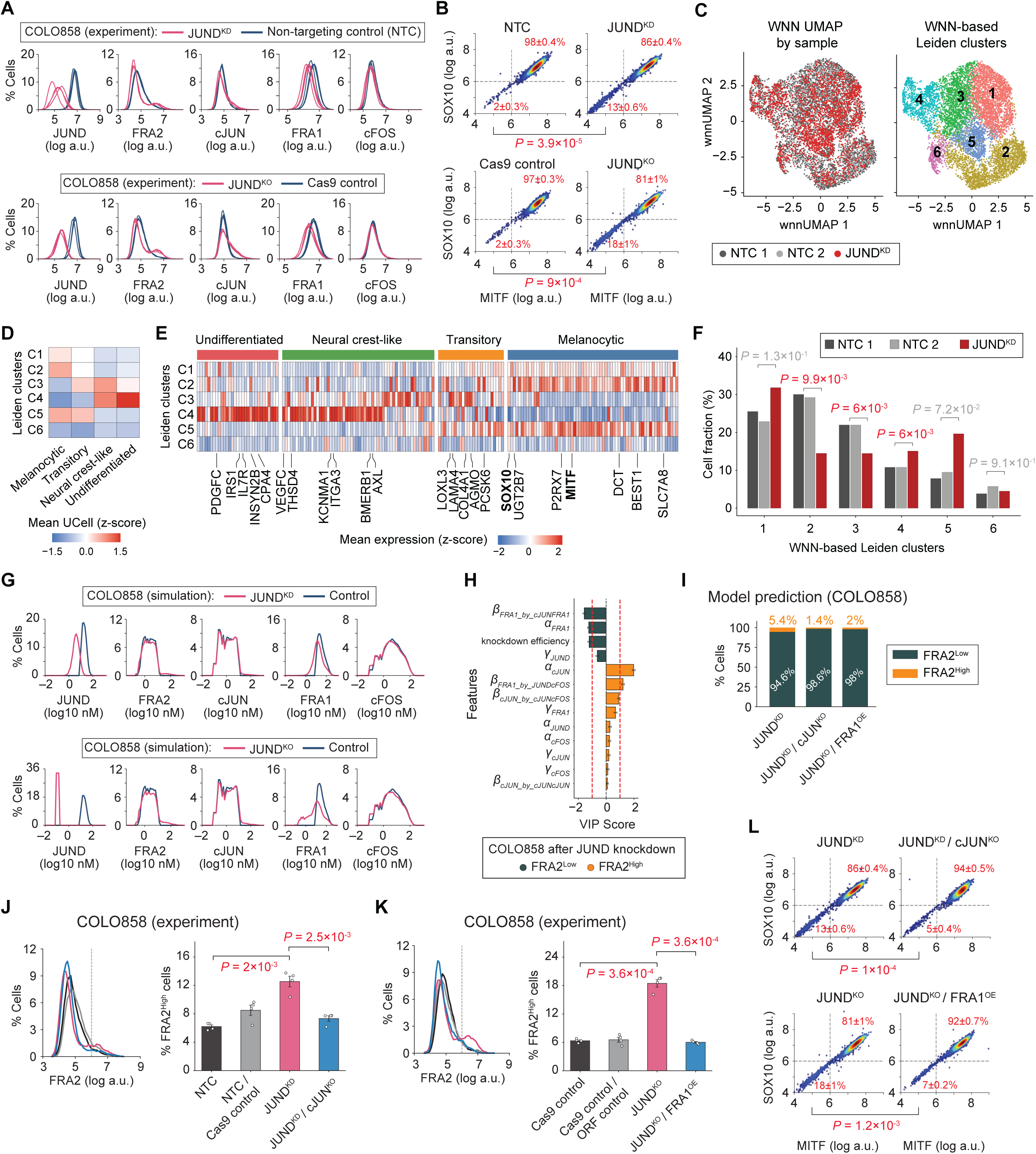
JUND loss promotes a FRA2^High^ state and drives an undifferentiated melanoma phenotype through reduced FRA1 and increased cJUN. **(A)** Single-cell distributions of AP-1 proteins (JUND, FRA2, cJUN, FRA1, and cFOS) measured by 4i across four replicates in COLO858 cells following shRNA-mediated JUND knockdown (top) or CRISPR/Cas9-mediated JUND knockout (bottom), compared with corresponding controls. **(B)** Single-cell covariance analysis of MITF and SOX10 protein levels measured by 4i across four replicates in COLO858 following JUND^KD^ or JUND^KO^ (right) and their respective controls (left). Dashed lines show thresholds defining undifferentiated (SOX10^Low^/ MITF^Low^) versus differentiated (SOX10^High^/ MITF^High^) subpopulations. Percentages of differentiated and undifferentiated cells (mean ± SEM across four replicates) are reported in the top-right and bottom-left quadrants, respectively. Statistical comparisons were performed using two-sided Welch’s *t*-tests. **(C)** Single-nucleus multiome (RNA + ATAC) analysis of COLO858 cells following JUND knockdown (JUND^KD^) and in control conditions (NTC 1 and NTC 2) using weighted nearest-neighbor (WNN) integration. UMAP plots show WNN embeddings colored by condition (JUND^KD^ vs NTC 1 vs NTC 2) (left) or by Leiden cluster assignment (right). **(D)** Population-averaged UCell enrichment scores for each of the four melanoma differentiation signatures (defined by Tsoi et al.) across Leiden clusters, z-scored across the six clusters. **(E)** Population-averaged, z-scored expression of melanoma differentiation genes across six Leiden clusters. Genes were selected based on their ability to distinguish clusters and their overlap with Tsoi et al. differentiation-state signatures. MITF and SOX10 (as key melanocyte lineage markers), and the top five genes per cluster are labeled. **(F)** Percentage of cells in each WNN-based Leiden cluster across JUND^KD^ and control (NTC 1 and NTC 2) samples. Benjamini-Hochberg adjusted *P* values assess differences in cluster proportions (from propeller) between JUND^KD^ and controls, using the two control replicates to estimate baseline variability. **(G)** Simulated single-cell distributions of the AP-1 proteins in virtual COLO858 populations following simulated JUND knockdown (top) or knockout (bottom), relative to the unperturbed control population. **(H)** Signed VIP scores from a PLS-DA model comparing simulated FRA2^High^ and FRA2^Low^ COLO858 subpopulations following virtual JUND knockdown, using AP-1 network parameters (excluding those directly regulating FRA2) as input features. **(I)** Model-predicted percentages of FRA2^High^ and FRA2^Low^ cells in COLO858 following combined perturbations: JUND knockdown (JUND^KD^) + cJUN knockout (cJUN^KO^), and JUND knockout (JUND^KO^) + FRA1 overexpression (FRA1^OE^). **(J-K)** Single-cell distributions of FRA2 protein levels measured by 4i across four replicates in COLO858 following JUND^KD^ alone or combined with cJUN^KO^ (J), and following JUND^KO^ alone or combined with FRA1^OE^ (K), compared to corresponding controls. Dashed lines indicate the FRA2 threshold used to identify FRA2^High^ cells. Statistical comparisons were made using two-sided Welch’s *t*-tests. **(L)** Single-cell covariance analysis of MITF and SOX10 protein levels measured by 4i across four replicates in COLO858 following JUND^KD^ alone (top left) or combined with cJUN^KO^ (top right), and following JUND^KO^ alone (bottom left) or combined with FRA1^OE^ (bottom right). Percentages of differentiated and undifferentiated cells (mean ± SEM across four replicates) are reported in the top-right and bottom-left quadrants, respectively. Statistical comparisons were performed using two-sided Welch’s *t*-tests.

To evaluate the impact of JUND depletion on differentiation-state heterogeneity beyond pre-selected markers, we performed single-cell multiome (RNA + ATAC) sequencing in COLO858 cells following JUND knockdown (JUND^KD^) and in non-targeting control (NTC) cells. To define cellular states, we constructed a joint RNA/ATAC embedding using the weighted nearest-neighbor (WNN) algorithm^55^. WNN-based Leiden clustering identified six subpopulations within the pooled COLO858 cell population (Figure 7C). Gene set enrichment analysis using the four melanoma differentiation-state signatures defined by Tsoi et al.^2^ associated four of these clusters with melanoma differentiation programs (Figures 7D and S7A). Cluster 4 (C4) was the only one enriched for the undifferentiated (U) signature and exhibited the lowest expression levels of both MITF and SOX10 (Figures 7D, 7E and S7B); cluster 2 (C2) was uniquely enriched for the melanocytic (M) signature; and clusters 5 (C5) and 3 (C3) displayed mixed melanocytic/transitory (M/T) and transitory/neural crest-like (T/N) signatures, respectively (Figures 7D, 7E).

To quantify the impact of JUND depletion on the distribution of these cell states, we applied propeller^56^, a linear modeling-based approach that leverages biological replication to find statistically significant differences in cell type proportions in single-cell data. Using two biological replicates for the control population (NTC 1 and NTC 2) to estimate baseline variability, we identified significant changes in the proportions of clusters C4, C2, and C3 following JUND knockdown that exceeded the variation observed between control replicates (Figure 7F). The undifferentiated cluster C4 expanded upon JUND depletion (*P* = 6×10^-3^), whereas the melanocytic cluster C2 showed the largest reduction (*P* = 9.9×10^-3^). In addition, the mixed-signature C3 population (transitory/neural crest-like) decreased in frequency (*P* = 6×10^-3^). Overall, these results indicate a significant redistribution of transcriptional cell states following JUND depletion. This shift is consistent with our 4i-based protein analysis, which independently demonstrated an increase in the fraction of MITF^Low^/SOX10^Low^ cells, supporting a transition toward an undifferentiated phenotype.

We next asked whether our computational model could predict the AP-1 reconfiguration underlying FRA2^High^ state induction. Simulating JUND depletion in a virtual population of COLO858 cells while capturing cell-to-cell heterogeneity, the model recapitulated the emergence of a small FRA2^High^ subpopulation accompanied by cJUN upregulation and FRA1 downregulation, consistent with experimental observations (Figure 7G). To identify molecular parameters associated with this transition, we constructed a PLS-DA model to distinguish FRA2^High^ and FRA2^Low^ cells following virtual JUND knockdown. AP-1 network parameters (excluding those directly regulating FRA2) were used as input features, and class imbalance was addressed through repeated random down-sampling. The model classified the two states with moderate accuracy (mean ROC AUC = 0.69 ± 0.02; Figure S7C). VIP analysis revealed that cells transitioning to the FRA2^High^ state had higher rates of cJUN production (either constitutive or induced by cJUN-cFOS dimer), whereas cells remaining FRA2^Low^ showed higher basal FRA1 production (α*_FRA1_*) or greater cJUN-FRA1 dimer-mediated induction of FRA1 (β*_FRA1_by_cJUNFRA1_*) (Figure 7H). This is consistent with our experimental observation that only a subset of FRA2^Low^ cell lines that downregulated FRA1 following ERK pathway inhibition exhibited FRA2 induction. To further evaluate the impact of α*_FRA1_* and β*_FRA1_by_cJUNFRA1_* on AP-1 network state under JUND depletion, we tracked model-predicted steady-state concentrations of AP-1 dimers across varying values of these parameters. Dose-response analysis showed that decreasing α*_FRA1_* and β*_FRA1_by_cJUNFRA1_*reduced levels of the cJUN-FRA1 heterodimer while increasing levels of cJUN-FRA2 and cJUN-cFOS dimers (Figure S7D). These shifts are consistent with remodeling of the AP-1 network toward enhanced cJUN and FRA2 production, mediated by cJUN-cFOS and cJUN-FRA2, respectively.

Consistent with these predictions, model simulations of either cJUN knockout (cJUN^KO^) or FRA1 overexpression (FRA1^OE^) in combination with JUND depletion reduced the fraction of FRA2^High^ cells compared to JUND^KD^ alone (Figure 7I). To validate these model predictions experimentally, we compared single-cell FRA2 protein expression in COLO858 cells following JUND^KD^ or JUND^KO^ alone and in combination with either cJUN knockout or FRA1 overexpression. Both perturbations significantly reduced the emergence of the FRA2^High^ subpopulation (Figures 7J and 7K), confirming that JUND loss promotes the FRA2^High^ state via a mechanism dependent on reduced FRA1 and increased cJUN.

Finally, given the strong association between the FRA2^High^ state and the undifferentiated phenotype, we hypothesized that cJUN^KO^ or FRA1^OE^ would also attenuate the shift toward an undifferentiated state observed following JUND depletion. To test this, we quantified SOX10 and MITF protein levels at single-cell resolution. The increased fraction of SOX10^Low^/ MITF^Low^ cells observed under JUND knockdown and knockout conditions was significantly reduced by both cJUN knockout and FRA1 overexpression (Figures 7L). Together, these results support a model in which AP-1 network remodeling downstream of ERK inhibition is associated with JUND loss, and JUND depletion is sufficient to induce an adaptive transition toward an undifferentiated phenotype. This transition is mediated by FRA1 downregulation and cJUN upregulation, promoting a FRA2^High^ state that is observed at a higher frequency in intrinsically undifferentiated melanoma tumors.

## Discussion

Cell state heterogeneity and plasticity are linked to metastasis and therapy resistance in melanoma and other cancers. Although MAPK signaling and AP-1 transcription factors have long been implicated in these processes, it has remained unclear how the AP-1 network generates distinct transcriptional states, why the distribution of those states differs across tumors and among cells within a tumor, and which upstream signals and network interactions control transitions between them. Answering these questions is essential for explaining the molecular origins of non-genetic heterogeneity and for designing strategies to modulate the state transitions that underlie tumor plasticity.

To address these gaps, we combined multiplexed single-cell protein measurements with mechanistic modeling. We identified six recurrent AP-1 expression states whose frequencies vary across 19 genetically distinct tumor lines and track their differentiation states. An ODE model encoding dimerization-mediated competition and dimer-specific auto- and cross-regulation captured the observed inter- and intra-line AP-1 heterogeneity and revealed parameter configurations that explain tumor line-specific AP-1 state distributions. Importantly, differences in inferred parameters correlated with MAPK pathway activities across lines and, within clonal populations, with the heterogeneity of these activities linking fluctuations in MAPK signaling to AP-1 states. Finally, we predicted and validated an adaptive AP-1 reconfiguration in which ERK inhibition induces JUND loss which, together with FRA1 downregulation and cJUN upregulation, promotes a FRA2^High^ state and a shift toward a dedifferentiated (SOX10^Low^/ MITF^Low^) phenotype, a state with well-known association with therapy resistance and pro-metastatic behavior^5,8,9^.

These findings support a network-level view of AP-1 as a configurable regulatory system whose phenotypic outputs are set by quantitative balances among competing AP-1 dimers and feedback interactions. In this view, JUND, maintained by ERK signaling, acts as a buffering node in differentiated melanoma cells; its loss following BRAF/MEK inhibition redistributes JUN partners, enabling FRA2 induction when FRA1 is concurrently reduced. The model also reconciles divergent roles previously attributed to individual AP-1 factors by showing that their effects depend on quantitative parameter regimes (including partner availability, production rates, and stability) that govern which dimers form and what outputs they drive. Because AP-1 controls plasticity programs across multiple cancers^21,57–59^, these principles likely extend beyond melanoma, with lineage- and mutation-specific differences determining which AP-1 components are most consequential. In melanoma, the role of JUND-FRA1-cJUN-FRA2 axis emerges as a key network mechanism of ERK inhibition-induced adaptive dedifferentiation and AP-1 driven therapeutic resistance.

Methodologically, this work illustrates a generalizable framework: integrating high-throughput multiplexed, single-cell measurements with mechanistic, single-cell models and statistical learning to infer regulatory parameters across diverse cellular contexts. Iterative integration of experiments and modeling enables the generation of testable hypotheses regarding the regulation of specific cell states and their responses to perturbations. This framework is extensible, e.g., with time-resolved calibration using live-cell reporters, or multiscale agent-based models that incorporate cell division, lineage history, and microenvironmental cues derived from spatial single-cell datasets. In parallel, emerging single-cell multiomic CRISPR screening platforms offer powerful approaches to identify core transcriptional programs underlying cell-state plasticity and therapy resistance-associated perturbations^60^. Given the central role of AP-1 network in integrating environmental signals, it will be also particularly important to map how AP-1 states are distributed within the tumor microenvironment, how paracrine cues shape AP-1 configurations, and how stromal and immune cells modulate or respond to AP-1 driven reprogramming. In this context, the high sparsity of AP-1 transcripts in single-cell RNA-seq datasets, together with their sensitivity to tissue dissociation, processing, and other *ex vivo* stressors inherent to these workflows, underscores the importance of *in situ* protein-level measurements. Multiplexed imaging approaches applied to intact tissues provide a particularly valuable strategy for resolving AP-1 network states and their contributions to tumor heterogeneity and plasticity.

### Limitations of the study

Several limitations of this study should be considered. First, our modeling framework is based on deterministic ODEs and emphasizes extrinsic variability in parameters and initial conditions as the primary driver of AP-1 state heterogeneity. While this approach defines stable states and their distributions, it does not explicitly capture intrinsic stochasticity in gene expression, which may also contribute to state transitions. Incorporating stochastic modeling approaches, together with future time-resolved single-cell measurements, will be important to quantify these effects. Second, model calibration relies on steady-state single-cell snapshots rather than dynamic measurements and uses discretized protein expression levels derived from fluorescence intensities in arbitrary units. In addition, AP-1 dimer concentrations are not directly measured but inferred through the model. Finally, although our study spans a broad panel of genetically and phenotypically diverse melanoma cell lines, the AP-1 protein measurements are performed *in vitro*, where the complexity of the tumor microenvironment is not fully captured. Given the central role of environmental signaling in shaping AP-1 activity, some AP-1 states observed in culture may differ from those in tumors.

## Methods

### Key resources table

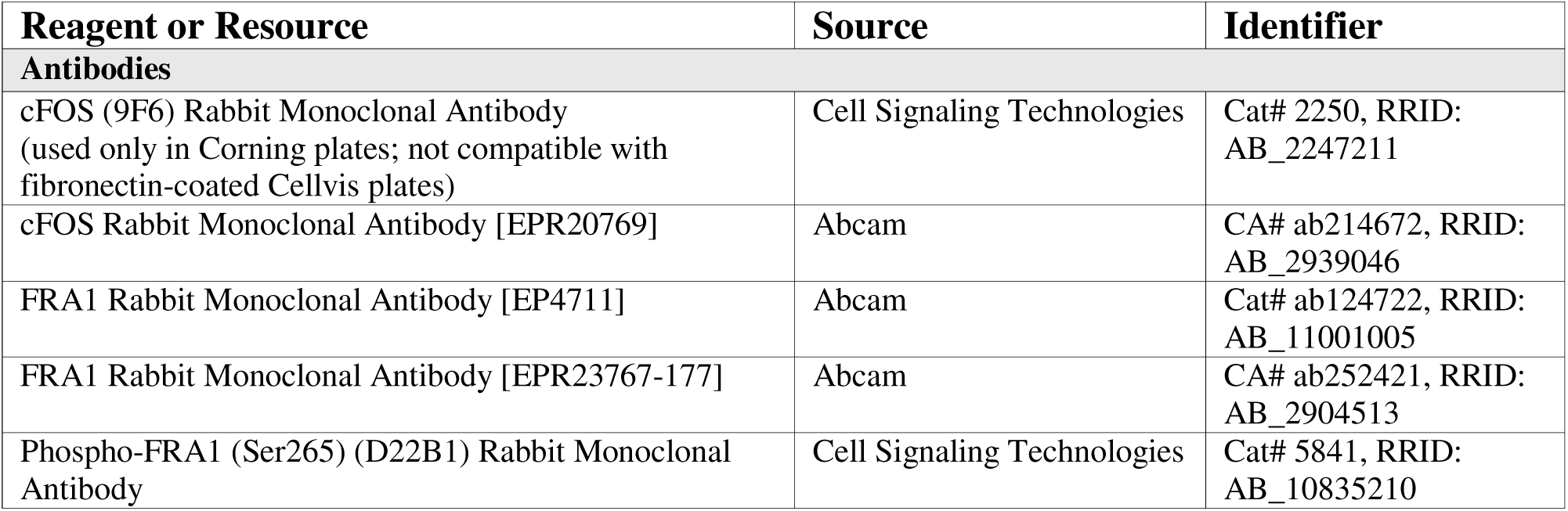
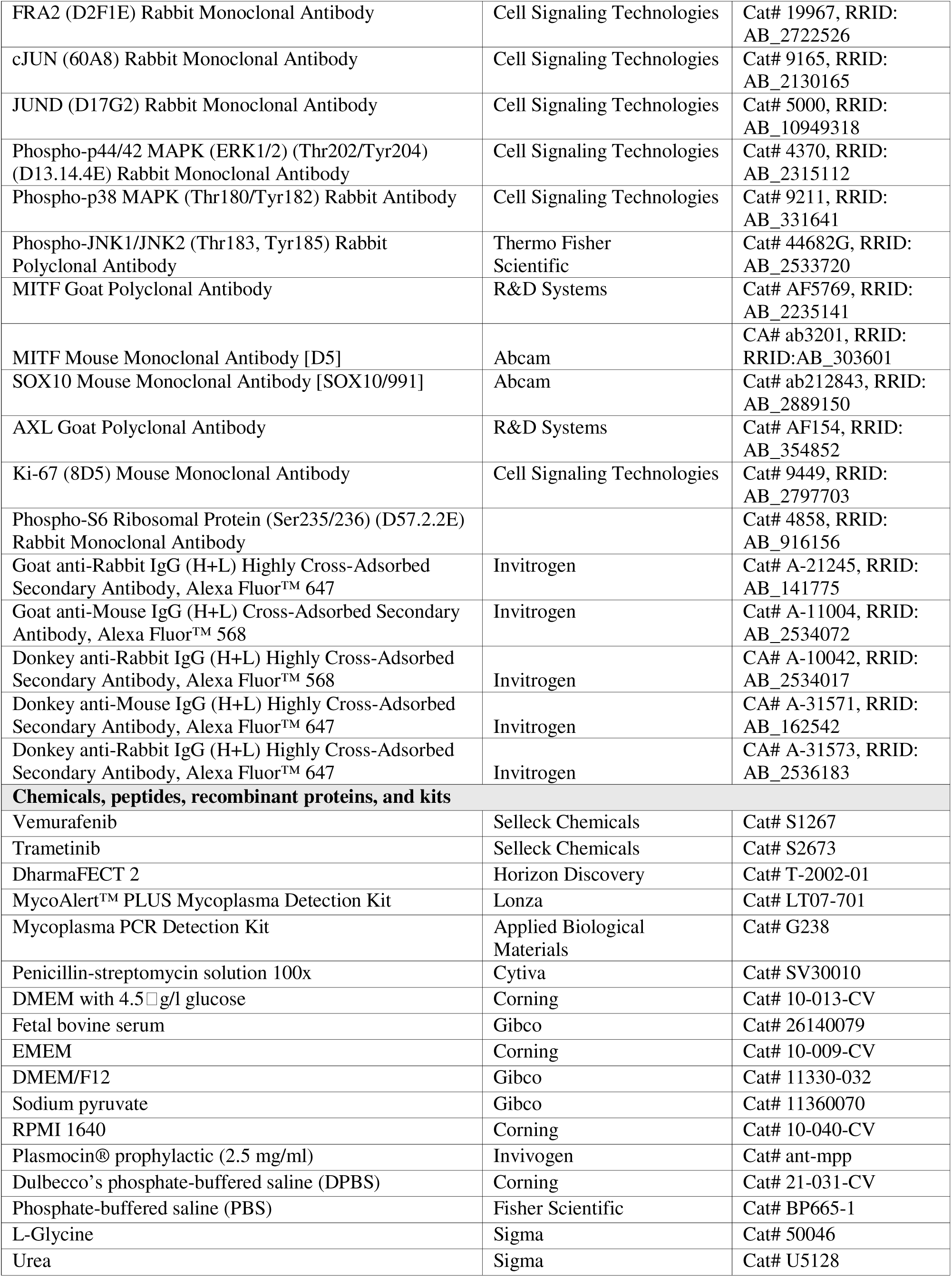
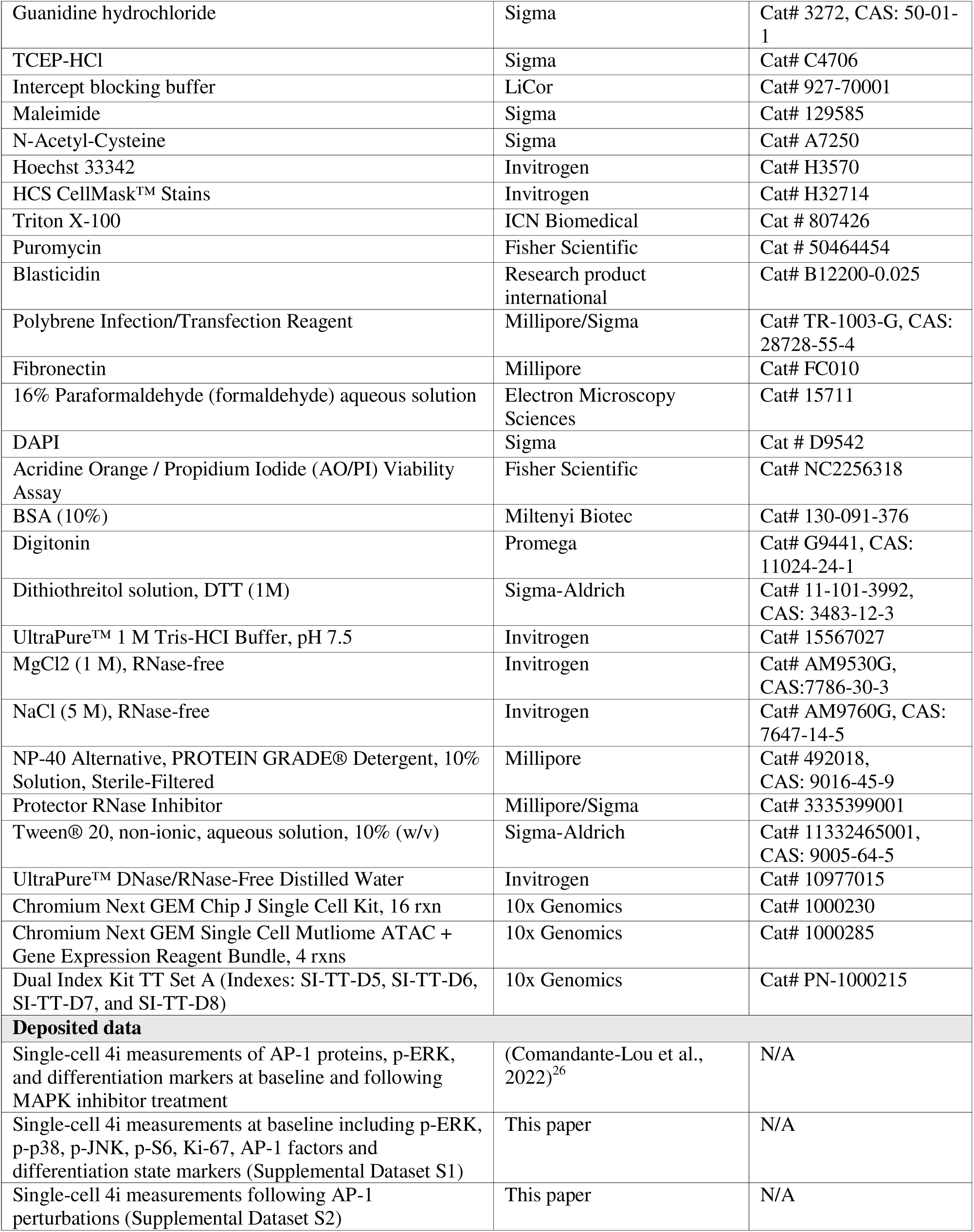
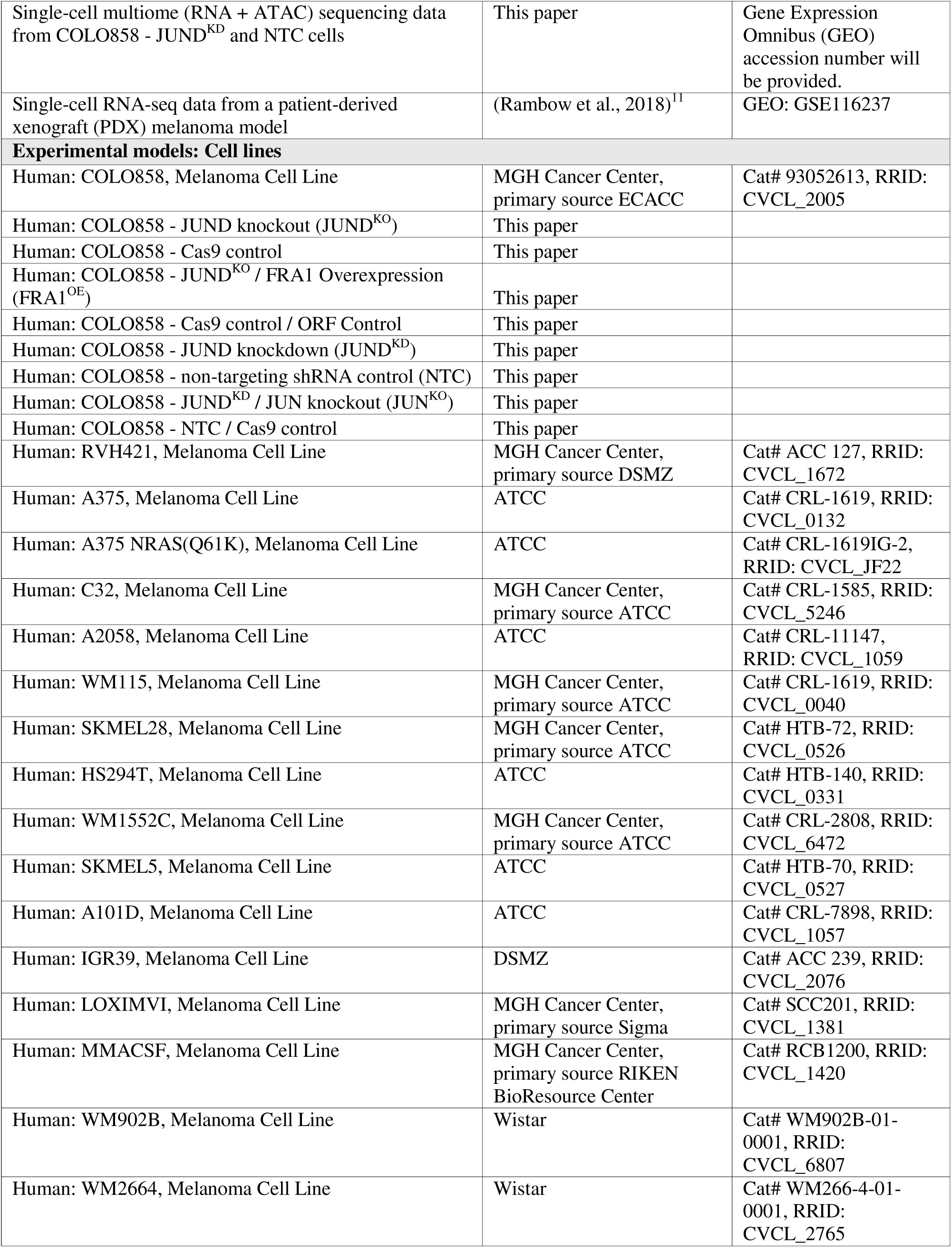
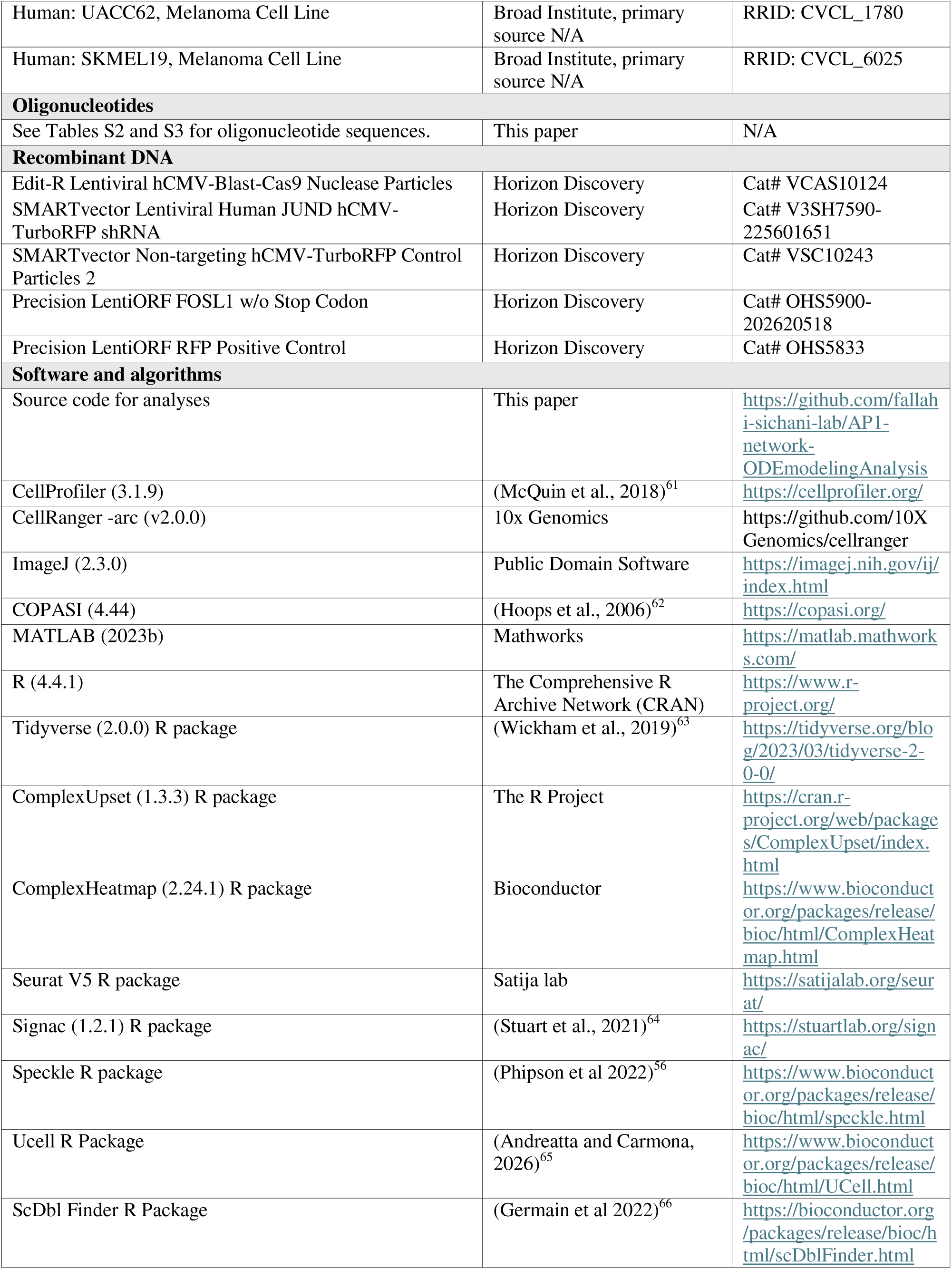
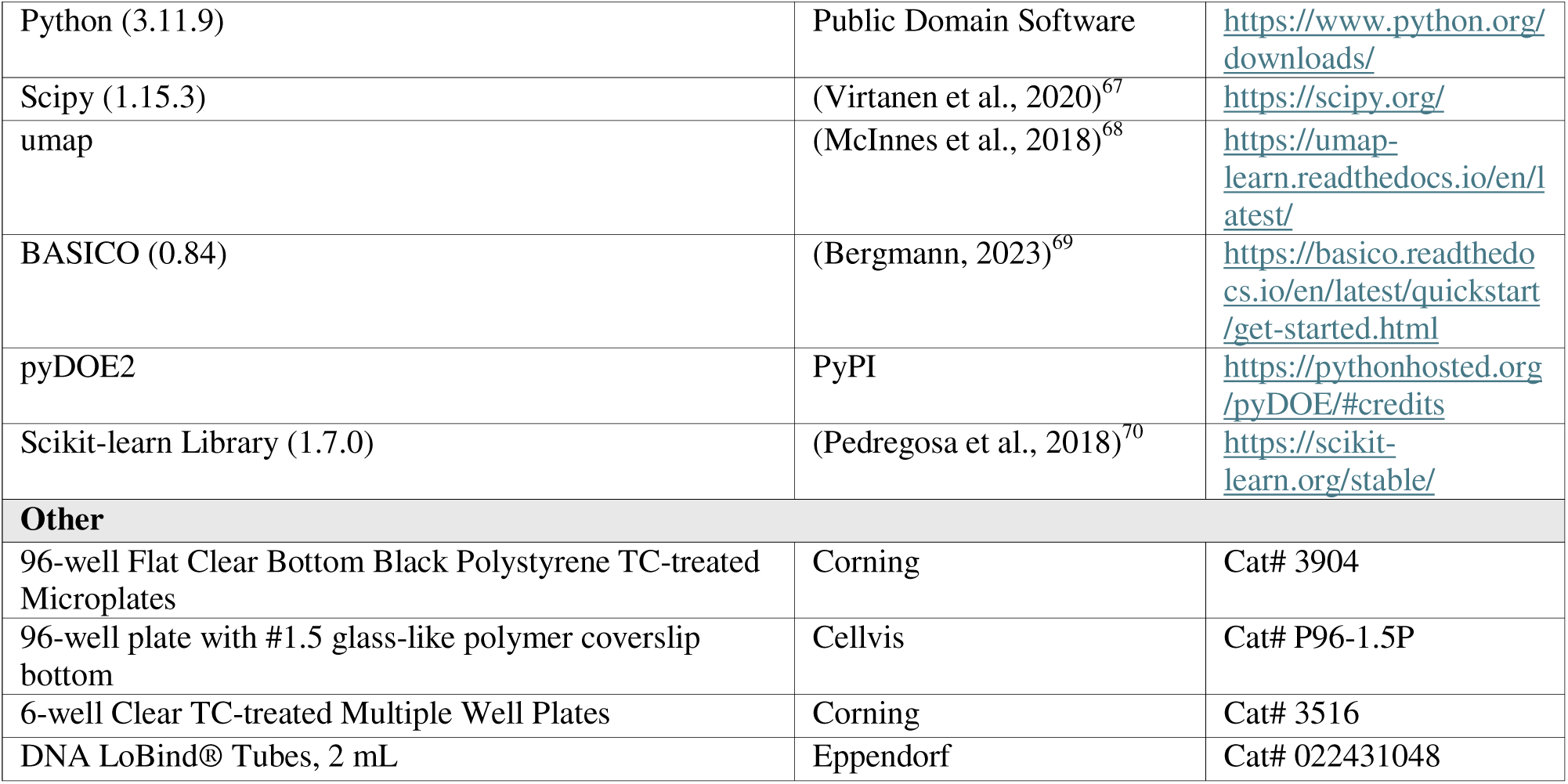

### Cell culture

BRAF-mutant melanoma cell lines used in this study include: COLO858, RVH421, A375, A375(NRAS^Q61K^), C32, A2058, WM115, SKMEL28, HS294T, WM1552C, SKMEL5, A101D, and IGR39, LOXIMVI, MMACSF, WM902B and WM2664, UACC62 and SKMEL19. All cell lines have been subjected to re-confirmation by short tandem repeat (STR) profiling by ATCC and mycoplasma testing by MycoAlert™ PLUS Mycoplasma Detection Kit or Mycoplasma PCR Detection Kit. A375, A375(NRAS^Q61K^), A2058, HS294T, A101D, and IGR39 cells were grown in DMEM with 4.5 g/l glucose supplemented with 5% fetal bovine serum (FBS). SKMEL5 and WM2664 cells were grown in EMEM supplemented with 5% FBS. C32, MMACSF, SKMEL28, and WM115 cells were grown in DMEM/F12 supplemented with 1% sodium pyruvate and 5% FBS. COLO858, LOXIMVI, RVH421, SKMEL19, UACC62, WM1552C, and WM902B cells were grown in RPMI 1640 supplemented with 1% sodium pyruvate and 5% FBS. 100 U/mL Penicillin-Streptomycin (10,000 U/mL) and 0.5 µg/mL Plasmocin Prophylactic were present in all cell cultures. COLO858 cells engineered by lentiviral transduction with vectors encoding either shRNA constructs or open reading frames (ORFs) were selected and thereafter maintained under antibiotic pressure: 0.5 µg/mL puromycin for shRNA-expressing lines and 5 µg/mL blasticidin for ORF-overexpressing lines. Matching control cell lines were also cultured under the same antibiotic conditions. Cells were grown at 37 °C with 5% CO_2_ in a humidified incubator.

### Iterative indirect immunofluorescence imaging (4i)

4i images were obtained using a cell culture-4i protocol previously described in full detail^71^. For the experiments performed on 19 melanoma cell lines (including drug treatments described below), cells were seeded in 200 µL/well in Corning 96-well plates in 2 biological replicates. For the experiments studying the impact of genetic perturbations (knockout, knockdown or overexpression) in AP-1 factors, cells were seeded in 200 μL/well in fibronectin-coated Cellvis 96-well plates at a density of 5,000 or 10,000 cells/well depending on their growth rate with 4 biological replicates each. Cells were imaged with a 10x air-objective on the Operetta CLS High-Content Imaging System (Revvity).

### Drug treatments

Drug treatments were done as described previously^26^. Cells were seeded in 200 µL/well in Corning 96-well plates. 0.316 µM Vemurafenib, 0.316 µM Vemurafenib + 0.0316 µM Trametinib or vehicle (DMSO) was added using the Tecan D300e Digital Dispenser 24 hours after cell seeding. Cells were fixed at the indicated timepoint (24 h) with 4% paraformaldehyde in phosphate-buffered saline (PBS) for 30 minutes at room temperature.

### 4i image analysis

The analysis of 4i data was done as previously described^26^. First, Image background subtraction was performed using the rolling ball subtraction algorithm in ImageJ (2.3.0) with a ball radius of 50. Background subtracted Hoechst images were aligned using the normalized cross correlation method within the Align module in CellProfiler (3.1.9)^61^. Nuclei were segmented from Hoechst channel images using the method that best captured cell line-specific nuclear morphology. For AP-1 perturbation experiments in COLO858, the Otsu method with three-class thresholding was used. Segmentation was performed using the “IdentifyPrimaryObjects” module in CellProfiler, assigning pixels in the middle intensity class to the foreground. The threshold smoothing scale and correction factor were set to 2.4 and 0.9 respectively. Cell segmentation was then performed based on images from the CellMask Green channel using Minimum Cross-Entropy thresholding method in the “IdentifySecondaryObjects” module. The threshold smoothing scale was set to 1 and the threshold correction factor was set to 0.5. Tracking of nuclei across all rounds of imaging was performed using Follow Neighbors method in the “TrackObjects” module. The mean intensity in the nuclei was extracted for all of the transcription factors, and the whole cell intensity was extracted for MAPK kinase activity (phosphorylation state) measurements. Additional organization and analysis of the data was performed with MATLAB.

### JUND and cJUN knockout by CRISPR-Cas9

COLO858 cells were seeded in a 6-well plate at a density of 150,000 cells/well in antibiotic-free growth medium. Two days post-seeding, cells were treated with 1 mL of transduction medium consisting of serum- and antibiotic-free growth medium supplemented with 10 µg/mL polybrene, and the Edit-R lentiviral hCMV-Blast-Cas9 nuclease particles (Horizon Discovery) at a multiplicity of infection (MOI) of 0.3. The plate was centrifuged at 2500 rpm for 30 minutes at 37°C and subsequently incubated at 5% CO_2_ at 37 °C in a humidified incubator. At 48 hours post-transduction, the medium was replaced with fresh growth medium supplemented with 5 µg/mL blasticidin, to enrich for Cas9-expressing cells. The concentration of blasticidin was determined using an antibiotic kill curve in COLO858 cells, in which 5 µg/mL of blasticidin was the minimum concentration that killed ∼100% of cells in less than 2 weeks. To generate cJUN and JUND knockouts, Cas9-expressing COLO858 cells were seeded in 6-well plates at a density of 120,000 cells/well in antibiotic-free medium. The next day, cells were transfected with 25 nM synthetic guide RNA complex (equimolar cJUN or JUND crRNA and tracrRNA) using 0.25 μL DharmaFECT 2 transfection reagent (Horizon Discovery) per well. Single-clone selection was performed using limited dilution (cJUN knockout) or single-cell sorting into 96-well plates based on ATTO 550 signal (with dead cell exclusion based on DAPI staining) using a Becton Dickinson Influx Cell Sorter (JUND knockout). Once clones were confluent, they were split into two 96-well plates. Protein depletion in selected cJUN and JUND knockout clones was verified by Western blotting and single-cell immunofluorescence imaging.

### JUND knockdown by lentiviral short hairpin RNA (shRNA)

To stably knock down JUND in parental COLO858 cells or in cJUN-knockout COLO858 cells, they were seeded in a 6-well plate at a density of 100,000 cells/well in antibiotic-free growth medium. Two days post-seeding, growth medium was aspirated, and cells were treated with 1 mL transduction medium consisting of serum- and antibiotic-free medium supplemented with 10 µg/mL polybrene and SMARTvector lentiviral human JUND hCMV-TurboRFP shRNA (Horizon Discovery) at MOI of 0.3. As control, parental COLO858 cells and Cas9-expressing COLO858 cells were also similarly transduced in parallel with SMARTvector non-targeting hCMV-TurboRFP control particles 2 (Horizon Discovery). After addition of transduction medium, the plate was centrifuged at 2500 rpm for 30 minutes at 37°C and subsequently incubated in a humidified incubator at 5% CO_2_ and 37 °C. 48 hours post-transduction, growth medium was replaced with fresh medium supplemented with 0.5 µg/mL puromycin, to enrich for the shRNA-expressing cells. The concentration of puromycin was determined using an antibiotic kill curve in COLO858 cells, in which 0.5 µg/mL of puromycin was the minimum concentration that killed ∼100% of cells in less than 1 week. To prevent shRNA-expressing cells from being outcompeted by cells that lost shRNA expression, they were continuously maintained in growth medium supplemented with puromycin at 0.5 µg/mL.

### FRA1 overexpression by lentiviral open reading frame (ORF)

To stably overexpress FRA1 in selected JUND-knockout COLO858 cells, cells were seeded in a 6-well plate at a density of 100,000 cells/well in antibiotic-free growth medium. Two days post-seeding, growth medium was aspirated, and cells were treated with 1 mL transduction medium consisting of serum- and antibiotic-free medium supplemented with 10 μg/mL polybrene, and Precision FOSL1 LentiORF w/o Stop Codon (Horizon Discovery) at a MOI of 0.3. As control, Cas9-expressing COLO858 cells were also similarly transduced in parallel with Precision LentiORF RFP Positive Control (Horizon Discovery). After the addition of transduction medium, the plate was centrifuged at 2500 rpm for 30 minutes at 37°C and subsequently incubated at 5% CO_2_ and 37°C in a humidified incubator. 48 hours post-transduction, growth medium was replaced with fresh medium supplemented with 5 μg/mL blasticidin and cells were maintained in this medium continuously. All ORF-positive cells were sorted based on their expression of TurboGFP using a Sony MA900 Cell Sorter.

### Isolation of nuclei and single-nucleus multiome library preparation

Nuclei for single-nucleus multiome analysis were isolated using the 10x Genomics® protocol for nuclei isolation from cell suspensions with minor modifications. COLO858 cells expressing non-targeting shRNA control (NTC; n = 2 biological replicates) or JUND knockdown (JUND^KD^) were seeded in T-75 flasks. Two days post-seeding, cells were washed with 10 mL of 1× DPBS and dissociated by incubation with 1 mL of 0.25% trypsin at 37°C for 30 seconds. Cell detachment was confirmed by microscopy, and cells were neutralized with 10 mL of growth medium. Cell suspensions were transferred to 15 mL tubes and centrifuged at 200 × g for 3 minutes at 4°C. The supernatant was removed, and cells were washed once with 5 mL of cold DPBS supplemented with 0.04% BSA, followed by centrifugation at 200 × g for 3 minutes at 4°C. The pellet was resuspended in 2 mL of cold DPBS with 0.04% BSA, and cells were counted. A total of 1×10^6^ cells were transferred to 2 mL LoBind tubes and centrifuged at 200 × g for 5 minutes at 4°C. The supernatant was carefully removed without disturbing the pellet. Cells were lysed by gentle resuspension in 100 µL of cold lysis buffer (0.1% NP-40, 0.1% Tween-20, 0.01% digitonin, 1% BSA, 1 mM DTT, 1 U/mL RNase inhibitor, 10 mM NaCl, 3 mM MgCl, 10 mM Tris-HCl, pH 7.5) and incubated on ice for 3 minutes. Nuclei were then washed three times with 1 mL of cold wash buffer (0.1% Tween-20, 1% BSA, 1 mM DTT, 1 U/mL RNase inhibitor, 10 mM NaCl, 3 mM MgCl, 10 mM Tris-HCl, pH 7.5). Following washes, nuclei were resuspended in 100 µL of 1× nuclei buffer supplemented with DTT and RNase inhibitor. Nuclei quality was assessed by acridine orange/propidium iodide (AO/PI) staining and imaging using a Leica Thunder Imaging Systems with a 40× air objective. Library preparation was performed by the University of Virginia Genome Analysis and Technology Core (RRID: SCR_018883) using the 10x Chromium Next GEM platform and Single Cell Multiome ATAC + Gene Expression reagents. Libraries were sequenced on a NextSeq 2000 platform.

### Single-nucleus RNA and ATAC multiome data processing and analysis

Raw BCL files were demultiplexed using *cellranger-arc* mkfastq, and per-library FASTQ files were aligned and quantified against the GRCh38 reference (refdata-cellranger-arc-GRCh38-2020-A-2.0.0) using *cellranger-arc* count (Cell Ranger ARC v2.0.0, default parameters). For the chromatin accessibility modality, peaks identified independently in each library by *cellranger-arc* were merged to generate a unified peak set across samples. ATAC fragments were then re-quantified against this common peak set using Signac^64^. Gene expression count matrices and re-quantified ATAC peak matrices were imported into R for downstream analysis.

Downstream analyses were performed in R using Seurat v5^72^ and Signac^64^. Nuclei were retained based on RNA quality-control criteria of 200-8,000 detected genes, ≤30,000 RNA UMIs, and ≤15% mitochondrial reads. Doublets identified using scDblFinder were excluded. Nuclei were further required to pass ATAC quality-control thresholds of 3,000-100,000 fragments, TSS enrichment >2, nucleosome signal <4, ENCODE hg38 blacklist ratio <0.05, and fraction of reads in peaks (FRiP) >0.15. Gene expression counts were normalized with SCTransform while regressing out percent mitochondrial reads and the difference between the S and G2M cell-cycle scores. Principal component analysis (PCA) was performed on SCTransform-normalized expression values, and the first 16 principal components were retained. ATAC peak counts were normalized using term frequency-inverse document frequency (TF-IDF) and reduced by latent semantic indexing (LSI). LSI components 2-25 were retained, excluding the first component due to its correlation with sequencing depth.

A joint RNA/ATAC embedding was constructed using Seurat’s weighted nearest-neighbor (WNN) algorithm, integrating the selected RNA principal components and ATAC LSI components. UMAP visualization and Leiden clustering were performed on the WNN graph at a resolution of 0.3. Per-sample cluster proportions were computed as the number of nuclei in each WNN cluster divided by the total nuclei in that sample. Differences in cluster abundance between NTC and JUND^KD^ conditions were assessed using *propeller*^56^ from the *speckle* package, based on logit-transformed cluster proportions, with *P* values adjusted using the Benjamini-Hochberg procedure. Variation between the two NTC biological replicates was used to estimate baseline variability; as JUND^KD^ was represented by a single replicate, the analysis assumes comparable variability between conditions.

To position nuclei along the melanoma differentiation trajectory, transcriptional signatures defined by Tsoi et al.^2^ were scored in each nucleus using UCell^65^, a rank-based signature scoring method on the RNA assay. To reduce the influence of sparsely detected genes, each signature was restricted to genes detected in at least 5% of nuclei. The seven Tsoi sub-signatures were consolidated into four canonical melanoma differentiation states, including melanocytic (M), transitory (T), neural crest-like (N), and undifferentiated (U) signatures, by taking the union of detection-filtered genes within each group^26^. To summarize differentiation-state enrichment across clusters, per-nucleus UCell scores were z-scored across nuclei and averaged within each cluster, and the resulting cluster-by-state matrix was visualized as a heatmap. UCell scores for each state were also projected onto the WNN UMAP as continuous color gradients. Expression of canonical melanocytic lineage markers (MITF and SOX10) was similarly visualized using SCTransform-normalized expression values. Cluster-specific marker genes were identified using Seurat’s *FindAllMarkers* function (Wilcoxon rank-sum test, one-versus-rest, positive markers only) applied to log-normalized RNA counts, with *P* values adjusted using the Benjamini-Hochberg procedure. To visualize differentiation-associated marker expression, genes meeting significance criteria (adjusted *P* < 0.05, log_2_-fold change > 0.5) and overlapping Tsoi differentiation signatures were selected, along with MITF and SOX10 as canonical markers. Expression values were averaged within clusters, z-scored across clusters, and grouped by differentiation state. All overlapping markers were displayed, with the top five genes per state (ranked by log_2_-fold change) highlighted.

### Single-cell RNA-seq analysis of patient-derived xenograft (PDX) models

We analyzed a publicly available melanoma PDX dataset in which single-cell RNA-seq was performed on melanoma cells isolated before treatment and following BRAF/MEK inhibitor therapy^11^. AP-1 transcript detection was assessed for FOS, FOSL1, FOSL2, JUN, and JUND by calculating, for each condition, the fraction of cells with non-zero expression for each gene. Due to high dropout rates across AP-1 genes (15-32% for FOS, 91-99% for FOSL1, 58-75% for FOSL2, 30-34% for JUN, and 17-20% for JUND), downstream analyses were restricted to univariate associations between individual AP-1 transcripts and Tsoi et al.^2^ melanoma differentiation-state signatures. For each AP-1 gene, Pearson correlation coefficients were computed between transcript expression and differentiation-state signature scores, as well as selected differentiation-associated markers, including NGFR and AXL. Correlations were calculated using only cells with detectable expression of the AP-1 transcript under consideration.

### Ordinary differential equation (ODE) modeling of the AP-1 network dynamics at a single-cell level

An ODE model was formulated to simulate the regulatory dynamics and competitive dimerization among five AP-1 transcription factors (cFOS, FRA1, FRA2, cJUN and JUND) in single cells based on mass action kinetics. The model includes AP-1 dimerization governed by protein-specific association (*k_on_*) and dissociation (*k_off_*) rate constants^27,30,41^. JUN subfamily proteins (cJUN, JUND) can form homodimers, while FOS family members (cFOS, FRA1, FRA2) require a JUN partner to dimerize. Each AP-1 protein (X) is produced at a basal rate (α*_X_*) representing its constitutive expression regulated by upstream signaling pathways. For proteins subject to auto- or cross-regulation, an additional dimer-induced production term (β*_X_by_YZ_*) was included to represent their regulated expression, modeled with a Hill function to capture cooperative dimer binding to AP-1 promoter sites^41^. Based on literature evidence, five dimer-mediated regulatory interactions were included: cJUN expression induced by cJUN-cJUN and cJUN-cFOS dimers; FRA1 induced by cJUN-FRA1 and JUND-cFOS; and FRA2 induced by cJUN-FRA2^30,40,42–46^. Four dimers not involved in AP-1 auto- or cross-regulation included JUND-JUND, JUND-cJUN, JUND-FRA1, JUND-FRA2^30,40,47,48^. Each protein (e.g., monomer X or dimer XY) degrades at a rate (γ*_X_* or γ*_XY_*, respectively) that depends on its identity and dimerization state^51,73–78^.

In addition, we made the following assumptions in model development and parameter estimation: (i) all reactions occur in a kinetically-homogeneous compartment in which all AP-1 protein concentrations are tracked until steady state at a single-cell level; (ii) AP-1 gene transcription and protein translation are lumped together and modeled as a single production event; (iii) we do not explicitly model AP-1 dimer binding to promoter regions. However, dimer-induced AP-1 production follows saturation kinetics (Hill function) with a half-maximal activation constant (*K_m_*) and cooperativity coefficient (*n*) that are equal for all transcriptionally active dimers^79,80^; (iv) Degradation rate constants for AP-1 dimers were computed as the arithmetic mean of the degradation rate constants of their constituent monomers, reflecting the idea that less stable AP-1 proteins gain stability through dimerization with more stable partners^75,81,82^; (v) JUN family proteins were estimated to have 100-fold higher affinity for forming heterodimers with FOS family members than for forming homodimers or heterodimers with other JUN proteins^42^.

We used the ODE model to track the concentrations of all AP-1 proteins (including monomers and dimers) at the single-cell level. To enable comparison with experimental data, we calculated the total concentration of each of the five AP-1 subunits (cFOS, cJUN, FRA1, FRA2, and JUND), regardless of their dimerization state. The construction of ODEs and computational simulations were performed using COPASI (version 4.38.30) and BASICO (version 0.51) for Python-COPASI interface. Python (version 3.11.9) was then used for data processing and visualization. Model parameters, their definitions, and their estimated ranges are included in Table S1. ODEs describing the dynamics of AP-1 proteins (e.g., monomer X, heterodimer XY, or homodimer XX) are listed below.

AP-1 monomers:

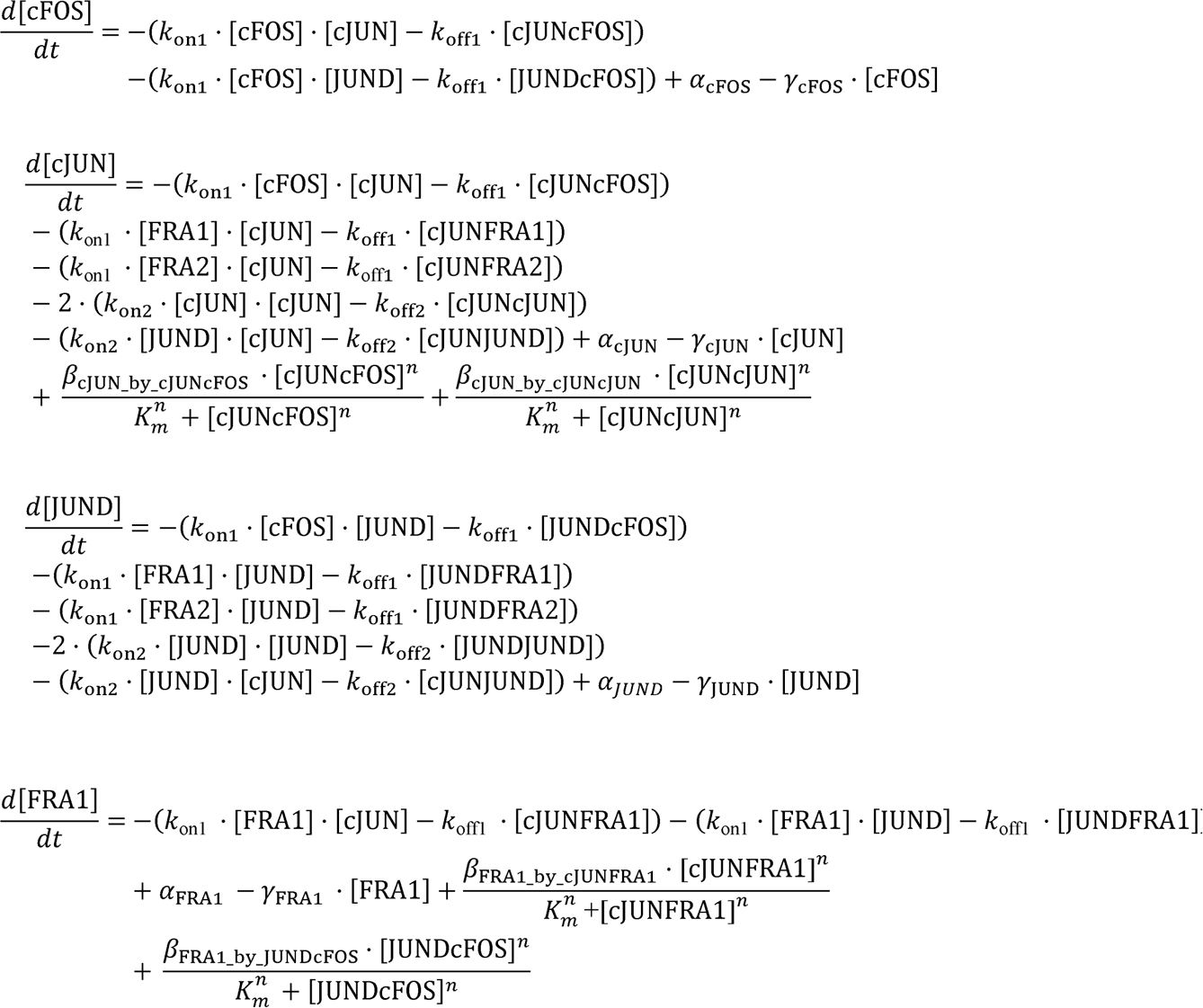

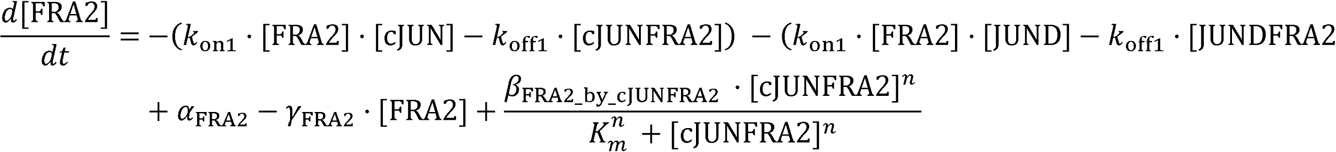

AP-1 dimers:

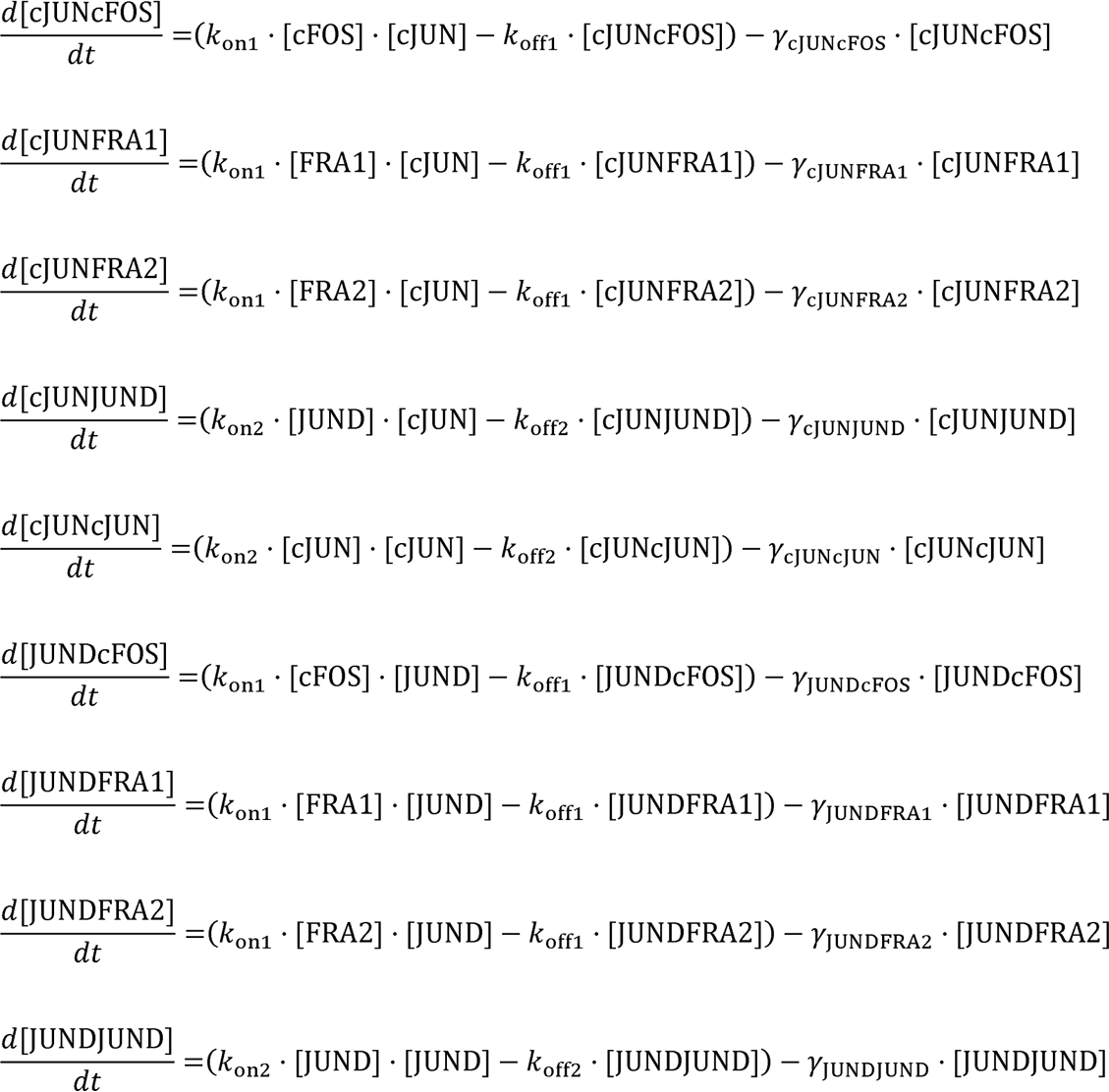

Total AP-1 concentrations:

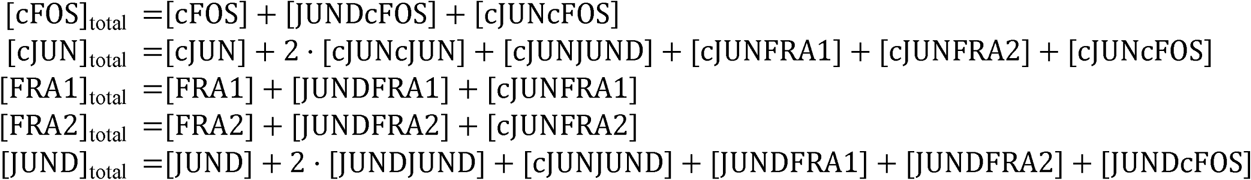

### ODE model parameter estimation and global scanning by Latin Hypercube Sampling (LHS)

Model parameters were either derived directly from the literature, or their physiologically relevant ranges were estimated according to the studies of similar transcription factors in mammalian cell systems, as described in Table S1. To generate single-cell heterogeneity in AP-1 states using ODE simulations, we incorporated extrinsic noise to capture cell type-specific variability. This included variation in the initial concentrations of all modeled AP-1 transcription factors (cFOS, FRA1, FRA2, cJUN, and JUND), as well as variability in kinetic rate parameters that may reflect cell-specific signaling activity. Specifically, we modeled extrinsic variability across 15 parameters, including basal production rates (α), dimer-induced production terms (β), and degradation rate constants (γ) for each AP-1 factor. For each parameter, we defined a physiologically relevant range based on the literature and constructed a 15-dimensional parameter space that was globally sampled and calibrated against single-cell data from 19 melanoma cell lines. In contrast, parameters governing AP-1 dimerization dynamics (*k_on_* and *k_off_* for homo- and heterodimers), the Hill coefficient (*n*) describing cooperative promoter binding, and the half-maximal activation constant (*K_m_*) for dimer-induced transcription were fixed based on published biochemical studies^27,30,41^, as these were assumed to be cell type-independent molecular properties. To simplify the model, we did not incorporate intrinsic noise arising from stochastic molecular interactions within the AP-1 network.

Cell type-specific parameter space was globally explored using Latin Hypercube Sampling (LHS), a computationally efficient approach for sampling multiple parameters simultaneously from a multidimensional distribution^49^. 20,000 parameter sets were randomly sampled using log-uniform distributions across physiologically relevant ranges defined for each parameter by using the lhs() function from the pyDOE2 library with random state fixed at 42, to ensure reproducibility. LHS was also used to sample 200 combinations of initial AP-1 protein concentrations (using log-uniform distributions) across a 1000-fold range, spanning from very low (0.316 nM) to very high (316 nM) levels for each protein. We then performed 4 million simulations by combining 200 initial conditions with 20,000 randomly sampled parameter sets. Steady state solutions were computed using the run steadystate() function in BASICO using the LSODA solver under default tolerance settings, and unique steady states across all simulations were determined using Kmeans clustering. About 2% of simulations failing to reach steady state due to numerical instability were excluded from subsequent analysis.

### Parameter calibration guided by experimentally recurrent AP-1 states

To identify regulatory configurations consistent with experimentally observed AP-1 state distributions across genetically distinct melanoma cell populations, we calibrated the 15-dimensional, cell type-specific parameter space to multiplexed single-cell data obtained from 19 melanoma cell lines.

Here, we defined an AP-1 state as the combinatorial expression pattern of five AP-1 proteins (cFOS, FRA1, FRA2, cJUN, and JUND) measured by 4i across these lines. To enable direct comparison between model predictions and experimental data (quantitative imaging in arbitrary units), total AP-1 protein levels in both datasets were discretized into binary “high” or “low” categories. This binarization allowed systematic identification of parameter sets compatible with the experimentally observed heterogeneity in each line. For model-predicted AP-1 levels, expression ≥10 nM was defined as high, and <10 nM as low. For 4i-derived AP-1 measurements across 19 cell lines, the following thresholds (in arbitrary units) were identified visually to classify individual melanoma cells as high or low expressers: cFOS (584 a.u.), cJUN (518 a.u.), FRA1 (518 a.u.), JUND (314.2 a.u.), and FRA2 (897.8 a.u.).

To identify the most robust AP-1 expression patterns observed experimentally across the 19 melanoma cell lines, we defined “recurrent states” as those present at an average frequency >10% (across two biological replicates) in at least three cell lines. To evaluate the robustness of AP-1 state classification to the choice of binarization thresholds, we performed a sensitivity analysis in which each AP-1 protein threshold was varied by ±0.15 natural log units, followed by re-identification of recurrent AP-1 states across all threshold combinations. In addition, we applied an alternative data-driven approach in which thresholds were estimated using two-component Gaussian mixture models fitted to the single-cell 4i data for each protein, followed by the same sensitivity analysis described above.

For cell line-specific calibration, all LHS-generated simulations were filtered to retain only combinations of parameter sets and initial conditions that reproduced the experimentally observed recurrent states. This filtering reduced the line-specific parameter subspace to experimentally consistent AP-1 states, captured either through retention (cells remaining in a given state) or transition (cells moving between two experimentally observed states). For LOXIMVI and COLO858, which were used in follow-up bistability analyses and AP-1 perturbation simulations, we performed an additional high-resolution LHS, generating 6 million simulations (40,000 parameter sets evaluated across 150 initial conditions) within cell line-specific parameter ranges, followed by calibration as described above.

To directly visualize the impact of key parameters (β*_cJUN_by_cJUNcJUN_* and β*_FRA2_by_cJUNFRA2_*) on FRA2 bistability in LOXIMVI cells, we generated bifurcation diagrams showing the steady-state concentration of total FRA2 or FRA2-containing dimers (JUND-FRA2 and cJUN-FRA2) as a function of these rate parameters in the LOXIMVI-calibrated model. For each parameter, values were varied over a 60-fold range on a logarithmic scale while all other parameters were held fixed at their calibrated values. At each parameter value, the model was initialized using the corresponding set of initial conditions and simulated to steady state. Simulations were classified as monostable when all initial conditions converged to a single steady-state solution, and as bistable when initial conditions converged to two distinct steady-state solutions.

### Simulating AP-1 perturbations in COLO858

We used combinations of parameter sets and initial conditions calibrated to COLO858 data to perform *in silico* (virtual) AP-1 knockdown, knockout, and overexpression experiments. To simulate JUND knockdown, while accounting for cell-to-cell heterogeneity in knockdown efficiency, we scaled the JUND basal production rate (α*_JUND_*) for each individual simulated cell (i.e., the *i*^th^ cell) by multiplying it by *k^(i)^*, the multiplier representing knockdown efficiency in that cell. *k^(i)^* values were drawn from a Beta distribution rescaled to the interval [0.01,1], corresponding to 0-99% knockdown efficiency. The shape parameters of the Beta distribution were tuned to ensure consistency between simulated and experimental JUND knockdown data, based on two comparison metrics: (i) the mean percent change of JUND relative to control cells following knockdown, and (ii) the overlap coefficient between single-cell JUND distributions in knockdown and control populations. To simulate JUND and cJUN knockouts, we set the corresponding production rate parameters (i.e., α*_JUND_* for JUND knockout and α*_cJUN_*, β*_cJUN_by_cJUNcJUN_*, and β*_cJUN_by_cJUNcFOS_*for cJUN knockout) to 0.001 (since setting their values to absolute zero would cause instability in numerical simulations) and verified that the knocked out protein would reach a concentration of ∼0 at steady state. FRA1 overexpression was modeled by increasing α*_FRA1_* 1000-fold relative to baseline. For combined perturbations, we applied knockouts first and then performed knockdown or overexpression starting from the knockout steady state, mirroring the order of these perturbations in the experimental work.

To evaluate the impact of α*_FRA1_* and β*_FRA1_by_cJUNFRA1_* on AP-1 network state under JUND depletion (JUND^KO^) in COLO858 cells, we performed a dose-response analysis by tracking model-predicted steady-state concentrations of AP-1 dimers across varying values of these parameters. In each analysis, either α*_FRA1_*or β*_FRA1_by_cJUNFRA1_*was varied over a defined range while all other parameters were held fixed at their calibrated values for the selected COLO858 parameter set. At each parameter value, the model was simulated to steady state, and the resulting AP-1 dimer concentrations were recorded.

### Partial least squares discriminant analysis (PLS-DA)

We used PLS-DA to identify molecular parameters associated with differences in AP-1 steady states simulated by ODE modeling, comparing them between different cell lines, between monostable and bistable outcomes in a single cell line (i.e., LOXIMVI), or between AP-1 states observed within a cell line. Calibrated parameters representing these conditions would serve as model input (i.e., predictive features). Parameter features used in PLS-DA were log-transformed and z-scored using StandardScaler () from the scikit-learn module. To address class imbalance in analyses where the majority class comprised more than 80% of samples, we applied conditional down-sampling to maintain a maximum 3:1 class ratio, using fixed random seeds for reproducibility. The optimal number of latent variables for PLS-DA analysis was determined using 5-fold stratified cross-validation with the StratifiedKFold() function, by maximizing the cross-validated area under the ROC curve (AUC). Model performance was assessed via cross-validation across 100 independent, random down-sampling iterations to account for variability arising from class imbalance, and mean AUC ± SD was reported. Variable importance in projection (VIP) scores were calculated using the optimal number of latent variables to rank the predictive features based on their contribution to model prediction. VIP scores were assigned directionality using logistic regression coefficients fitted on the same preprocessed data, where positive signed VIP scores indicated positive correlation with the positive class while negative signed VIP scores indicated negative correlation with the positive class. VIP > 1 or VIP < -1 indicated significant contribution to the class prediction by the model. For models with class imbalance, VIP scores were computed across down-sampling iterations and reported as mean ± SD.

### Dimensionality reduction by principal component analysis (PCA) and uniform manifold approximation and projection (UMAP)

PCA was performed using python’s scikit-learn module following z-scoring the data. UMAP was performed using the umap package in python. To visualize single-cell protein measurements across all cell lines, we first performed PCA using python’s scikit-learn module on z-scored log-transformed data and retained PCA scores from the first three principal components to use for the UMAP analysis. Parameters used for the UMAP analysis include nearest neighbor (neighbors = 90), minimum distance (min_dist) = 0.7, and distance metric (metric) = Euclidean. To overlay simulated AP-1 steady states with experimental single-cell data in a shared UMAP space, we independently z-scored the log-transformed simulation and experimental datasets. For each cell line, the experimental data were randomly subsampled (without replacement) to match the number of cells in the corresponding simulated dataset, ensuring equal sample sizes per cell line. The combined dataset was then visualized by UMAP with the following parameters: neighbors = 50, min_dist = 0.5, metric = Euclidean.

### Hierarchical clustering

We used hierarchical clustering using the stats package in R (4.4.1) to analyze expression patterns of the five AP-1 transcription factors (cFOS, cJUN, FRA1, FRA2, JUND) and the differentiation markers (MITF, SOX10). Single cell protein expression data from 19 melanoma cell lines were aggregated by calculating mean expression level for each protein within each cell line. Protein expression values were z-scored within each protein across cell lines. Clustering was performed using the hclust () function using the average algorithm as the agglomeration method, and Pearson correlation as the distance metric using the dist() function.

### Quantification and statistical analysis

Single-cell protein abundance was quantified from microscopy images using CellProfiler (3.1.9). No statistical method was used to predetermine sample size. Sample sizes were chosen based on similar studies in the relevant literature. The experiments were not randomized. The investigators were not blinded to allocation during experiments and outcome assessment. All violin plots highlight the median, lower and upper quartiles. Sample sizes (i.e., number of cells or replicates) are indicated in the figures. The statistical significance of differences between experimental or simulated single-cell datasets (sampled in equal sizes), either across cell lines or between subpopulations within a cell line, was assessed using two-sided Wilcoxon rank-sum tests. For the single-cell level evaluation of which cell lines exhibited JUND or FRA1 down-regulation following drug treatments, a one-sided Wilcoxon sign-ranked test was used to determine the *P* value. Statistical significance for the difference between the single cell distribution of PLS-DA scores (across latent variables) was evaluated using a Kolmogrov-Smironv test (KS test). Comparison between COLO858 cells following genetic perturbations of AP-1 factors was performed using Welch’s two sample *t*-test using randomly sampled equal number of cells from four replicates. Statistical analyses were performed using Python (version 3.11.9).

## Resource availability

### Lead contact

Further information and request for resources should be directed to and will be filled by the lead contact, Mohammad Fallahi-Sichani.

### Materials availability

This study did not generate new unique reagents.

### Data and code availability

- Raw experimental data reported in this paper will be shared by the lead contact upon request. The quantified single-cell microscopy data are included in Supplemental Datasets S1 and S2. This paper analyzes existing, publicly available data. The information for the public datasets are listed in the key resources table.
- The original codes for data analysis performed in this paper are publicly available at GitHub: https://github.com/fallahi-sichani-lab/AP1-network-ODEmodelingAnalysis
- Any additional information required to reanalyze the data reported in this paper is available from the lead contact upon request.

## Acknowledgments

We thank members of the Fallahi-Sichani laboratory for their technical contributions, helpful suggestions, and discussion. We acknowledge the University of Virginia (UVA) Flow Cytometry Core Facility (RRID: SCR_017829), Genome Analysis and Technology Core (RRID: SCR_018883), and UVA Research Computing for providing technical support and computational resources. This work was supported by NIH grants R35-GM133404, T32-GM145443, T32-GM007267, T32-CA009109, and P30-CA044579 (University of Virginia Cancer Center Support Grant). The Sony MA900 cell sorter used in this study was funded through the NIH S10 instrumentation program (S10-OD028518).

## Author Contributions

Y.N.D. and M.F.-S. conceived and designed the study and wrote the manuscript. Y.N.D. performed the computational modeling work. D.G.B. and M.B. performed the experiments. Y.N.D. and M.B. and D.G.B. analyzed the experimental data. M.F.-S. supervised the work.

## State of Competing Interests

The authors declare that they have no competing interests.

## Declaration of generative AI and AI-assisted technologies in the manuscript preparation process

During the preparation of this work, the authors used Claude and ChatGPT to assist with coding, debugging, and refactoring of the computational analysis code. After using these tools, the authors reviewed and edited the content as needed and take full responsibility for the content of the published article.

