## Supplemental Information for "Cell state plasticity emerging from co-regulated, competitive, and configurable interactions within the AP-1 network"

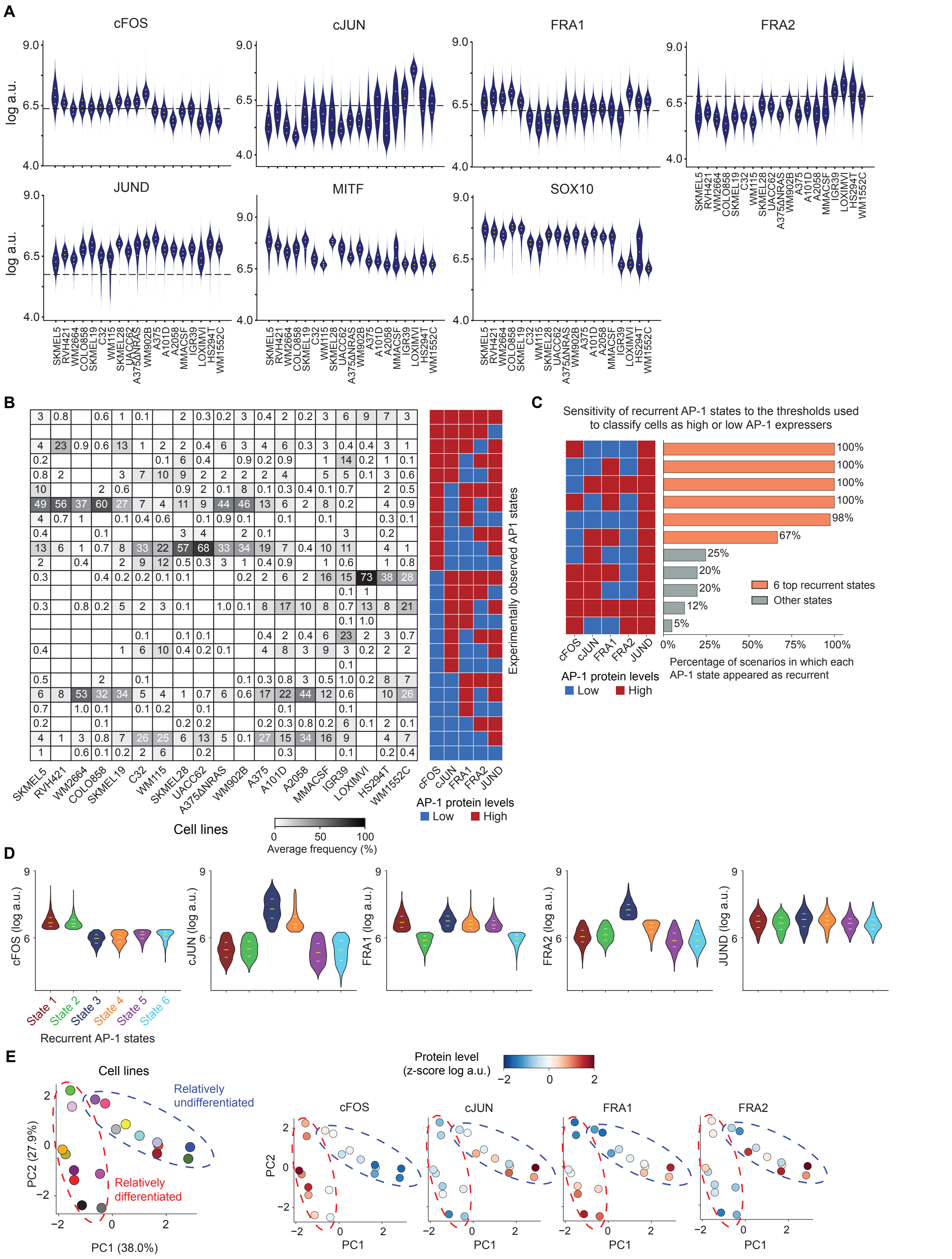

**Figure S1. Melanoma cells express six recurrent AP-1 states linked to differentiation states across genetically diverse populations. (A)** Single-cell distributions of five AP-1 factors (cFOS, cJUN, FRA1, FRA2, JUND) and two differentiation state markers (MITF and SOX10) measured across 19 cell lines and shown by violin plots highlighting the median and interquartile (25%

and 75%) ranges. **(B)** Frequency of cells in each of the 24 experimentally observed AP-1 states (defined by the combinatorial expression of five AP-1 proteins) across 19 melanoma cell lines. Data are averaged across two biological replicates. **(C)** Sensitivity of recurrent AP-1 states to threshold selection. Recurrent AP-1 states were defined as those present at an average frequency >10% (across two biological replicates) in at least three of the 19 melanoma cell lines. To assess the robustness of state classification, binarization thresholds for each AP-1 protein were varied by ±0.15 natural log units, and recurrent states were re-identified across all threshold combinations, and percentage of scenarios in which each AP-1 state appeared as recurrent was shown. **(D)** Single-cell distributions of five AP-1 protein levels among the six recurrent AP-1 states combined across all cell lines. Violin plots show median and interquartile ranges. **(E)** Principal component analysis (PCA) of melanoma cell lines based on their single-cell frequencies in the six recurrent AP-1 states. Cell lines are colored either by identity (left) or by population-averaged expression of AP-1 proteins (right).

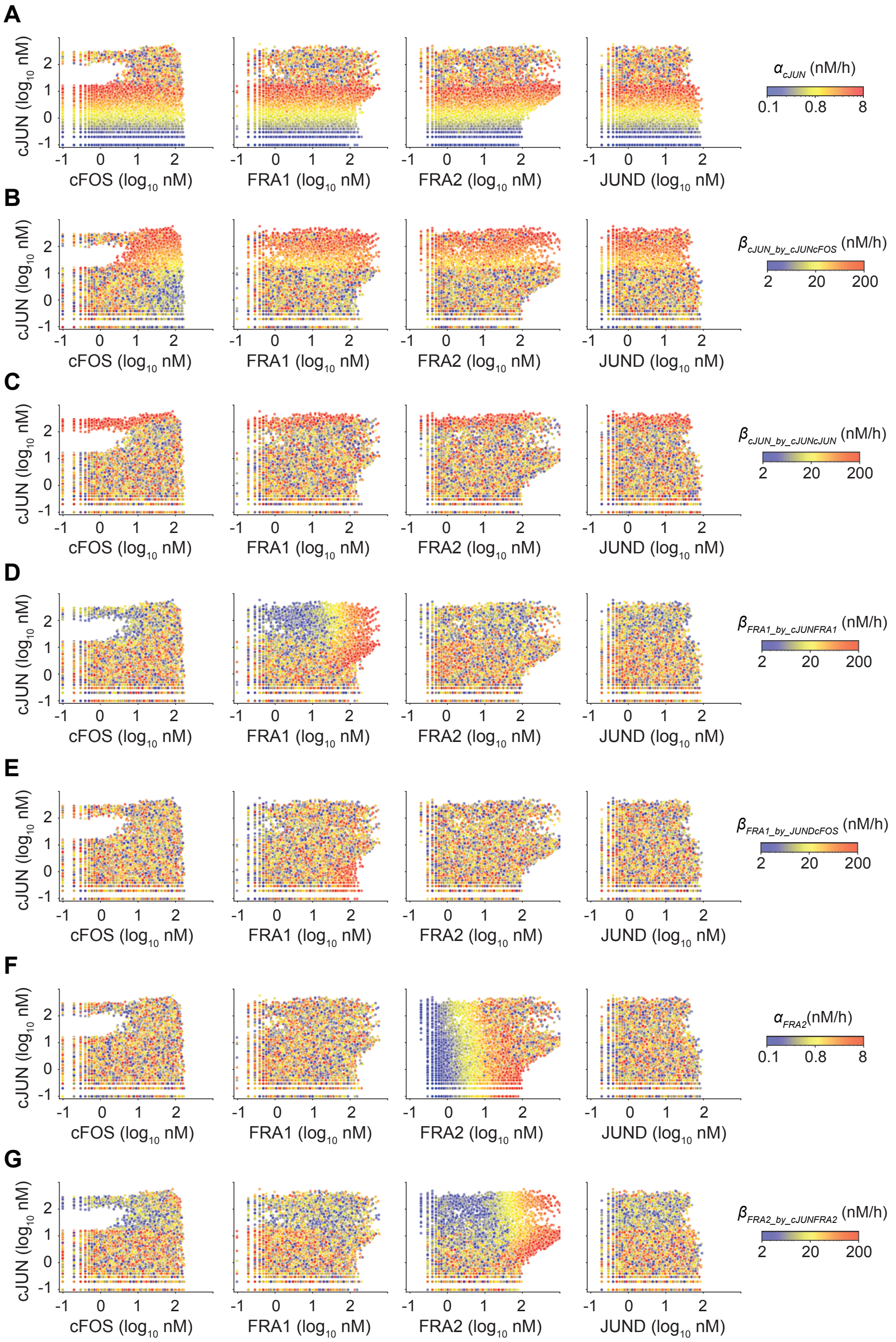


**Figure S2. Visualization of AP-1 steady-state outcomes as a function of individual parameters.**
Steady state outcomes from 4 million dynamic simulations, combining 200 initial conditions with 20,000 parameter sets sampled by Latin Hypercube Sampling (LHS), are shown. **(A-C)** Effects of basal production (A) or dimer-induced production of cJUN mediated by cJUN-cFOS homodimer (B) or cJUN-cJUN heterodimer (C). **(D, E)** Effects of dimer-induced production of FRA1 mediated by cJUN-FRA1 dimer (D) or JUND-cFOS dimer(E). **(F, G)** Effects of basal (F) or dimer-induced production of FRA2 mediated by cJUN-FRA2 dimer (G).

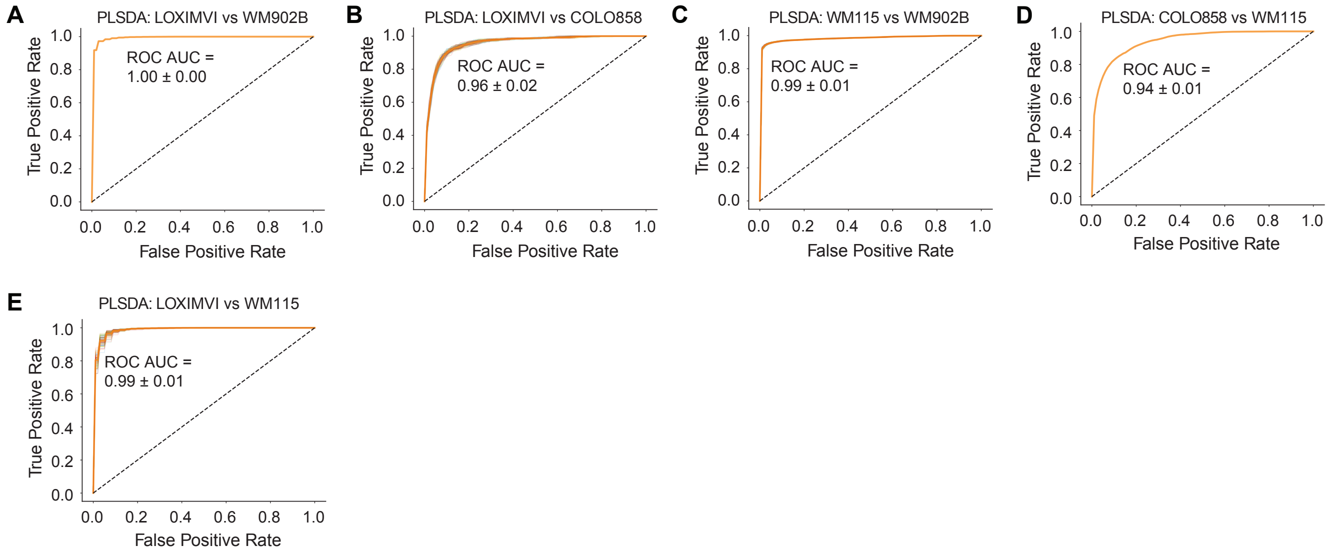


**Figure S3. The area under the ROC curve shown as an overall measure of the PLS-DA classifier performance for pairwise comparisons across cell lines. (A)** PLS-DA model comparing cell populations from LOXIMVI versus WM902B. **(B)** PLS-DA model comparing cell populations from LOXIMVI versus COLO858. **(C)** PLS-DA model comparing cell populations from WM115 versus WM902B. **(D)** PLS-DA model comparing cell populations from COLO858 versus WM115. **(E)** PLS-DA model comparing cell populations from LOXIMVI versus WM115. Model performance was evaluated by five-fold stratified cross-validation, with the area under the ROC curve (AUC) reported as mean ± SD.

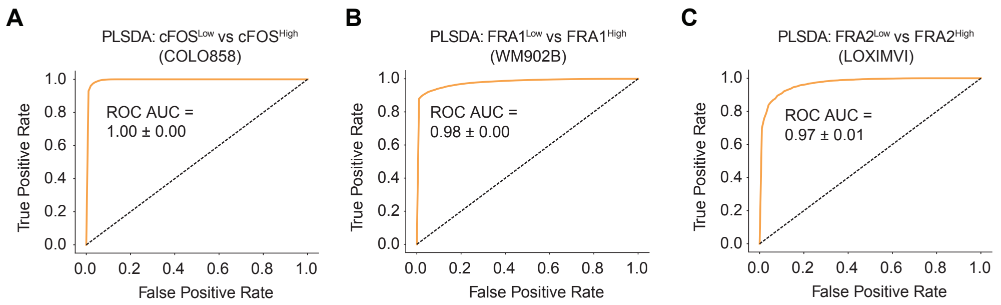


**Figure S4. The area under the ROC curve shown as an overall measure of the PLS-DA classifier performance for comparison of cell subpopulations within cell lines. (A)** PLS-DA model comparing $cFOS^{High}$ and $cFOS^{Low}$ subpopulations in COLO858. **(B)** PLS-DA model comparing $FRA1^{High}$ and $FRA1^{Low}$ subpopulations in WM902B. **(C)** PLS-DA model comparing $FRA2^{High}$ and $FRA2^{Low}$ subpopulations in LOXIMVI. Model performance was evaluated by five-fold stratified cross-validation, with the area under the ROC curve (AUC) reported as mean ± SD.

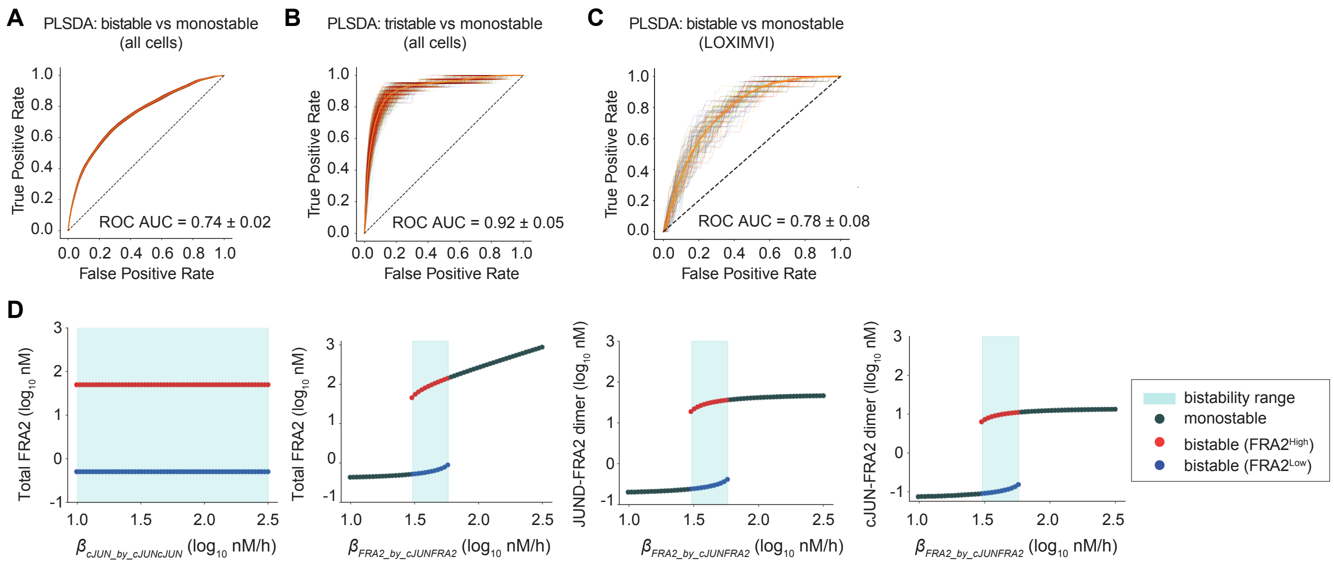


**Figure S5. The balance between basal and dimer-induced FRA2 production determines FRA2 expression bistability. (A-C)** The area under the ROC curve shown as an overall measure of the PLS-DA classifier comparing bistable and monostable regimes among all cells (A), comparing tristable and monostable regimes among all cells (B), and comparing bistable and monostable regimes in LOXIMVI. Model performance was evaluated using five-fold stratified cross-validation, and the area under the ROC curve (AUC) is reported as mean ± SD. **(D)** Bifurcation diagrams showing the steady-state concentration of total FRA2 or FRA2-containing dimers (JUND-FRA2 and cJUN-FRA2) as a function of $\beta_{cJUN\_by\_cJUNcJUN}$ and $\beta_{FRA2\_by\_cJUNFRA2}$ for LOXIMVI cells. For each parameter, values were varied over a 60-fold range on a logarithmic scale while all other parameters were held fixed at their calibrated values. At each parameter value, the model was initialized using the corresponding set of initial conditions and simulated to steady state. Simulations were classified as monostable when all initial conditions converged to a single steady-state solution, and as bistable when initial conditions converged to two distinct steady-state solutions.

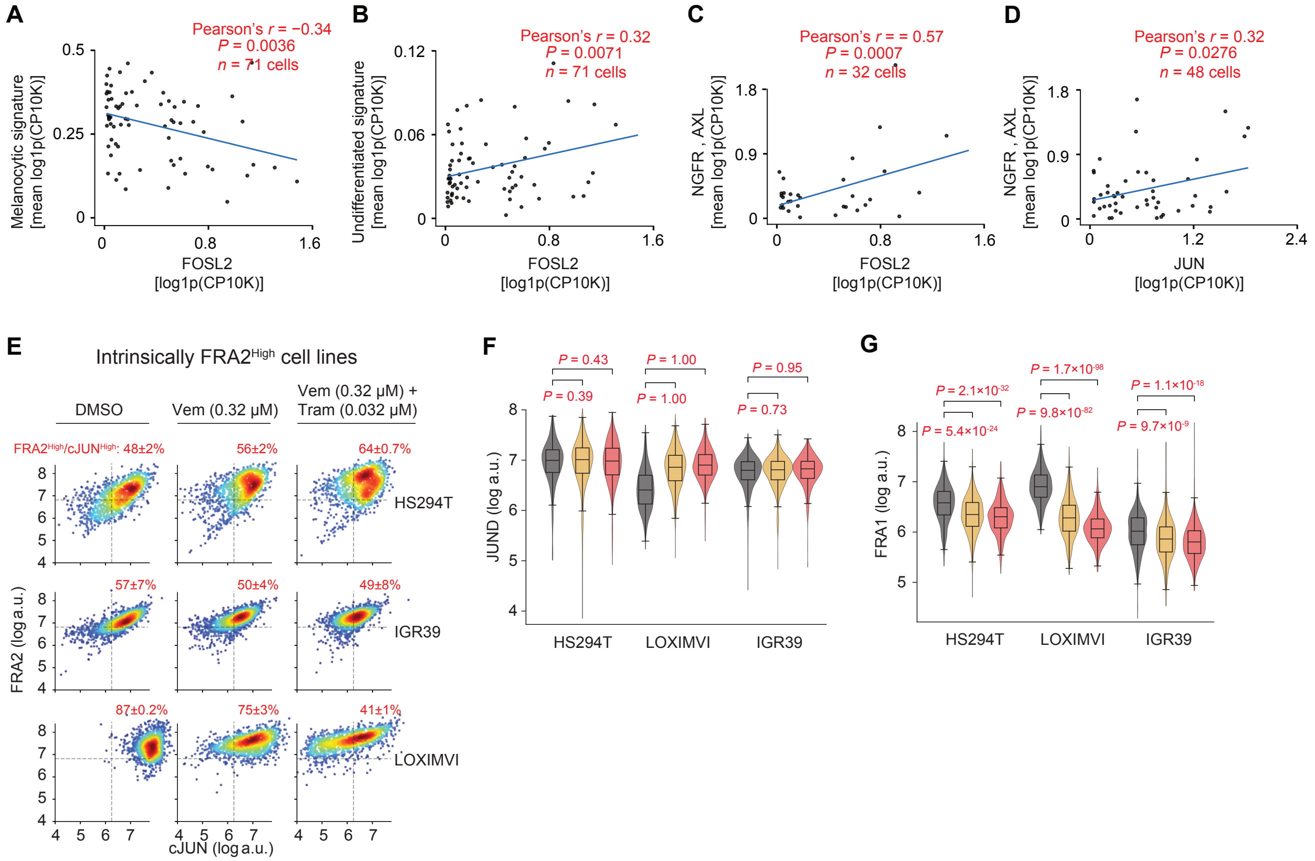


**Figure S6. MAPK inhibitor-induced FRA2 upregulation is accompanied by cJUN upregulation, JUND downregulation, and FRA1 downregulation. (A-C)** Univariate associations between FOSL2 transcript abundance and the Tsoi et al. melanocytic gene signature (A), undifferentiated gene signature (B), and undifferentiated melanoma markers NGFR+AXL (C). **(D)** Univariate association between JUN transcript abundance and undifferentiated melanoma markers (NGFR+AXL). Data are derived from single-cell RNA-seq analysis of a melanoma PDX dataset generated by Rambow et al. Correlations were calculated using only cells with detectable expression of the AP-1 transcript under consideration. **(E)** Single-cell covariance analysis of FRA2 and cJUN protein levels in melanoma cells following 24 h treatment with Vemurafenib (0.32 µM), the combination of Vemurafenib (0.32 µM) and Trametinib (0.032 µM), or vehicle control (DMSO), measured by 4i. Data are shown for intrinsically $cJUN^{High}$/ $FRA2^{High}$ cell lines. The percentage of $cJUN^{High}$/ $FRA2^{High}$ cells (averaged across two replicates ± SD) is shown in the top-right quadrant of each plot. **(F,G)** Single-cell analysis of JUND (F) and FRA1 protein levels (G) in melanoma cells following 24 h treatment with Vemurafenib (0.32 µM), the combination of Vemurafenib (0.32 µM) and Trametinib (0.032 µM), or DMSO, measured by 4i. Data are shown for intrinsically $cJUN^{High}$/ $FRA2^{High}$ cell lines. Statistical comparisons were performed using one-sided Wilcoxon signed-rank tests. Box plots display medians and interquartile ranges.

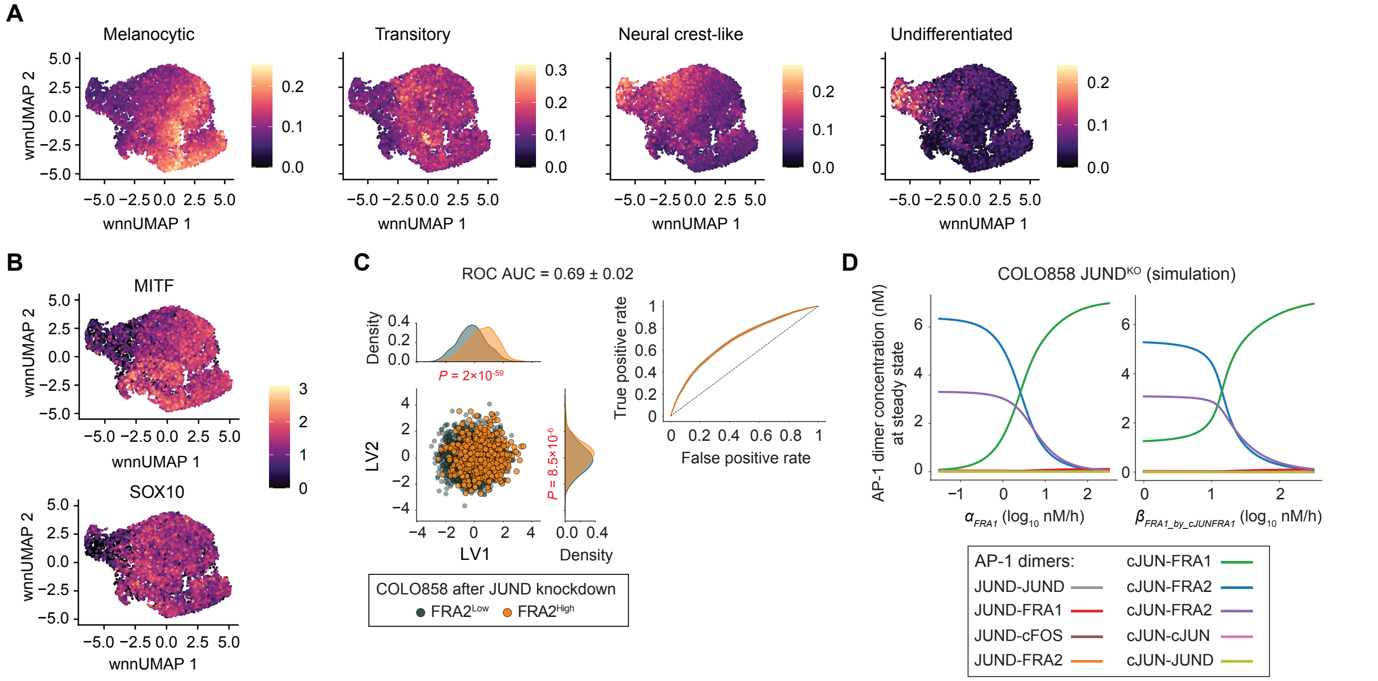


**Figure S7. JUND loss promotes a FRA2$^{High}$ state and drives an undifferentiated melanoma phenotype through reduced FRA1 and increased cJUN. (A,B)** Single-nucleus multiome (RNA + ATAC) analysis of COLO858 cells following JUND knockdown (JUND$^{KD}$) and in control conditions (NTC 1 and NTC 2) using weighted nearest-neighbor (WNN) integration. UMAP plots show WNN embeddings colored by Tsoi et al. differentiation-state signatures, including melanocytic, transitory, neural crest-like and undifferentiated states (A) or expression levels of MITF and SOX10 (B). **(C)** PLS-DA model comparing simulated FRA2$^{High}$ and FRA2$^{Low}$ COLO858 subpopulations following virtual JUND knockdown, using AP-1 network parameters (excluding those directly regulating FRA2) as input features. The PLS-DA score plot (left) shows individual cells colored by FRA2 expression state, with distributions compared across latent variables. Statistical significance along each latent variable was assessed using the Kolmogorov-Smirnov (KS) test. Model performance was evaluated using five-fold stratified cross-validation, and mean ± SD area under the ROC curve (AUC) is shown (right). **(D)** The impact of $\alpha_{FRA1}$ (left) and $\beta_{FRA1\_by\_cJUNFRA1}$ (right) on AP-1 network state under JUND depletion (JUND$^{KO}$) in COLO858 cells, assessed by a dose-response analysis via tracking model-predicted steady-state concentrations of AP-1 dimers across varying values of these parameters.

**Table S1. Model parameters, their description, and estimated values or ranges used in this study.**

| Parameter | Description | Unit | Value or Range | Source |
|---|---|---|---|---|
| $\alpha_{cFOS}$ | Basal production rate of cFOS | nM/h | 0.08 - 80 | Zhu et al., 2022 |
| $\alpha_{cJUN}$ | Basal production rate of cJUN | nM/h | 0.08 - 8 | Zhu et al., 2022 |
| $\alpha_{FRA1}$ | Basal production rate of FRA1 | nM/h | 0.08 - 8 | Zhu et al., 2022 |
| $\alpha_{FRA2}$ | Basal production rate of FRA2 | nM/h | 0.08 - 8 | Zhu et al., 2022 |
| $\alpha_{JUND}$ | Basal production rate of JUND | nM/h | 0.08 - 8 | Zhu et al., 2022 |
| $\beta_{cJUN\_by\_cJUNcFOS}$ | Production rate of cJUN induced by cJUN-cFOS dimer | nM/h | 2 - 200 | Zhu et al., 2022 |
| $\beta_{FRA1\_by\_cJUNFRA1}$ | Production rate of FRA1 induced by cJUN-FRA1 dimer | nM/h | 2 - 200 | Zhu et al., 2022 |
| $\beta_{FRA2\_by\_cJUNFRA2}$ | Production rate of FRA2 induced by cJUN-FRA2 dimer | nM/h | 2 - 200 | Zhu et al., 2022 |
| $\beta_{cJUN\_by\_cJUNcJUN}$ | Production rate of cJUN induced by cJUN-cJUN dimer | nM/h | 2 - 200 | Zhu et al., 2022 |
| $\beta_{FRA1\_by\_JUNDcFOS}$ | Production rate of FRA1 induced by JUND-cFOS dimer | nM/h | 2 - 200 | Zhu et al., 2022 |
| $\gamma_{cFOS}$ | Degradation rate of cFOS monomer | 1/h | 0.426 - 1.704 | Basbous et al., 2007; Shah and Tiyagi, 2013 |
| $\gamma_{cJUN}$ | Degradation rate of cJUN monomer | 1/h | 0.417 - 1.668 | Hernandez et al., 2008; Shah and Tiyagi, 2013 |
| $\gamma_{FRA1}$ | Degradation rate of FRA1 monomer | 1/h | 0.174 - 0.694 | Basbous et al., 2007; Casalino et al., 2003 |
| $\gamma_{FRA2}$ | Degradation rate of FRA2 monomer | 1/h | 0.8 - 0.32 | Alli et al., 2013 |
| $\gamma_{JUND}$ | Degradation rate of JUND monomer | 1/h | 0.058 - 0.232 | Hernandez et al., 2008 |
| $\gamma_{cJUNcFOS}$ | Degradation rate of cJUN-cFOS dimer | 1/h | $(\gamma_{cJUN} + \gamma_{cFOS}) / 2$ | Estimated |
| $\gamma_{cJUNFRA1}$ | Degradation rate of cJUN-FRA1 dimer | 1/h | $(\gamma_{cJUN} + \gamma_{FRA1}) / 2$ | Estimated |
| $\gamma_{cJUNFRA2}$ | Degradation rate of cJUN-FRA2 dimer | 1/h | $(\gamma_{cJUN} + \gamma_{FRA2}) / 2$ | Estimated |
| $\gamma_{cJUNJUND}$ | Degradation rate of cJUN-JUND dimer | 1/h | $(\gamma_{cJUN} + \gamma_{JUND}) / 2$ | Estimated |
| $\gamma_{cJUNcJUN}$ | Degradation rate of cJUN-cJUN dimer | 1/h | $\gamma_{cJUN}$ | Estimated |
| $\gamma_{JUNDJUND}$ | Degradation rate of JUND-JUND dimer | 1/h | $\gamma_{JUND}$ | Estimated |
| $\gamma_{JUNDcFOS}$ | Degradation rate of JUND-cFOS dimer | 1/h | $(\gamma_{JUND} + \gamma_{cFOS}) / 2$ | Estimated |
| $\gamma_{JUNDFRA1}$ | Degradation rate of JUND-FRA1 dimer | 1/h | $(\gamma_{JUND} + \gamma_{FRA1}) / 2$ | Estimated |
| $\gamma_{JUNDFRA2}$ | Degradation rate of JUND-FRA2 dimer | 1/h | $(\gamma_{JUND} + \gamma_{FRA2}) / 2$ | Estimated |
| $k_{off1}$ | Dimer dissociation rate between JUN and FOS family members | 1/h | 14.4 | Kohler and Shepartz, 2001 |
| $KD_1$ | Dimer binding affinity between JUN and FOS family members | nM | 2.8 | Reinke et al., 2013 |
| $k_{on2}$ | Dimer association rate between JUN family memebers | 1/(nM.h) | $k_{off2}$ / $KD2$ | Derived |

| | | | | |
|---|---|---|---|---|
| $k_{off2}$ | Dimer dissociation rate between JUN family memebers | 1/h | 1444 | Kohler and Shepartz, 2001 |
| $KD_2$ | Dimer binding affinity between JUN family members | nM | 100 × *KD1* | Estimated; Grondin et al., 2007 |
| $k_{on1}$ | Dimer association rate between JUN and FOS family members | 1/(nM.h) | $k_{off1}$ / $KD_1$ | Derived |
| *n* | Hill Coefficient used to capture cooperative dimer binding to AP-1 promoter sites | dimensionless | 1.5 | Bintu et al., 2005; Ajo-Franklin et al., 2007 |
| $K_m$ | half-maximal activation constant for dimer-induced AP-1 production | nM | 10 | Bintu et al., 2005; Ajo-Franklin et al., 2007 |

**Table S2. Edit-R synthetic crRNA and tracrRNA used for generating knockout cell lines.**

| Oligonucleotide | Source, Cat# | Target sequence |
|---|---|---|
| **Edit-R Human JUN (3725) crRNA** | Horizon Discovery, Cat# CM-003268-01-0002 | GTTGAGGGCATCGTCATAGA |
| **Edit-R Human JUND (3727) crRNA** | Horizon Discovery, Cat# CM-003900-01-0002 | TAGAGGAACTGTGAGCTCGT |
| **Edit-R CRISPR-Cas9 Synthetic tracrRNA** | Horizon Discovery, Cat#, U-002005-05 | |
| **Alt-R™ CRISPR-Cas9 tracrRNA, ATTO™ 550** | IDT, Cat# 1075927 | |

**Table S3. Primers used for validation of knockout cell lines.**

| Target gene | Forward primer | Reverse primer |
|---|---|---|
| **JUN** | GAGGTGAGGAGGTCCGAGTT | GCGTGCGCTCTTAGAGAAAC |
| **JUND** | CGGGAAGGGCACAGGTT | CGCCTCATCATCCAGTCCAA |
